# A Comparison and Validation of BIDS Apps for Preprocessing of [^11^C]UCB-J PET data

**DOI:** 10.64898/2026.10.28.762181

**Authors:** Guusje Haver, Granville James Matheson, Hampus Yngwe, Max Andersson, Niklas K Edvall, Martin Nørgaard, Francesca Zanderigo, Mike Schmidt, Elizabeth A Bartlett, Jeffrey M Miller, Zsolt Cselényi, Mikael Tiger, Maria Beckman, Carl-Johan Ekman, Johan Lundberg

**Affiliations:** Centre for Psychiatry Research, Department of Clinical Neuroscience, Karolinska Institutet & Stockholm Health Care Services, Region Stockholm, Sweden; Department of Psychiatry, Vagelos College of Physicians and Surgeons at Columbia University, New York, NY, USA; Department of Computer Science, University of Copenhagen, Copenhagen, Denmark; Molecular Imaging Branch, National Institute of Mental Health (NIMH), United States; Pathophysiology and Treatment Research Division, New York State Psychiatric Institute, New York, NY, USA; PET Science Centre, R&D, AstraZeneca, Stockholm, Sweden

**Keywords:** Positron Emission Tomography, BIDS, preprocessing, synaptic density, reproducibility

## Abstract

**Background:** Preprocessing of Positron Emission Tomography (PET) data can be performed in many different ways and often involves manual steps, resulting in reduced comparability across analyses and pipelines. Apps working on PET data organized according to the Brain Imaging Data Structure (BIDS) seek to automate preprocessing, enabling the application of standardized pipelines that increase comparability and reproducibility. In this study, two of these apps were validated against in-house software.

**Methods:** Data were acquired in-house from 14 subjects who received placebo in a randomized controlled clinical trial and underwent two PET measurements using the radioligand [^11^C]UCB-J. Preprocessing was performed with two BIDS Apps, PETPrep and PICNIC, as well as an in-house software. Arterial blood data were processed using Bloodstream, and [^11^C]UCB-J total distribution volume (*V*_T_) values were calculated using a 1-tissue compartment model using kinfitr.

**Results:** *V*_T_ values for all 9 regions of interest were highly consistent across tools with estimates of Intraclass Correlation Coefficients for consistency ranging between 0.94 and >0.99, although bias between tools measured as signed Percentage Difference was present, ranging from 0.1% – 10.2% depending on the region. Tools performed similarly on test-retest metrics. We observed a negative test-retest bias in [^11^C]UCB-J *V*_T_ of unknown origin.

**Conclusion:** BIDS Apps produce reliable and consistent results when compared both with each other and with an in-house software optimized for processing and analysis of our in-house data. Region-dependant bias between tools warrants caution for direct comparison of *V*_T_ between tools. The usage of these apps could enable better comparability across publications.

ClinicalTrials.gov NCT04630964 (https://clinicaltrials.gov/study/NCT04630964)

## Introduction

Processing and analysis of Positron Emission Tomography (PET) data is often performed using in-house software or closed-source commercial tools, which introduces an unknown degree of variability between centers at each processing step. Moreover, even using open-source software, there are many ways in which different preprocessing steps and choices can be combined into pipelines^1^, and these steps are often insufficiently described in a given scientific article to be fully reproduced^2^. The NeuroReceptor Mapping (NRM) Grand Challenge in 2018 highlighted how variability in (pre)processing strategies used by different PET research groups around the world propagated into variability in downstream sensitivity to detect changes in simulated PET data^3^.

The Brain Imaging Data Structure (BIDS) is a community-driven standard for organisation, storage and sharing of neuroimaging data, which emphasises both machine- and human-readability^4^. BIDS was originally developed for Magnetic Resonance Imaging (MRI), but has been expanded to an increasing number of modalities in the field of neuroimaging^5^, with PET added in 2022^6^. PET-BIDS is now the *de facto* community standard for sharing PET data^7^, such as through OpenNeuroPET^8^. Besides data sharing, by virtue of its standardised structure and machine-readability, BIDS also enables the development and application of fully automated processing pipelines, called BIDS Apps^9^. BIDS Apps are tools that ingest data stored in BIDS format and perform various preprocessing or analysis steps. The advantage of BIDS Apps over other preprocessing software is that they can be applied to data from any research group provided it is stored in the BIDS format, and these tools can be deployed without local installations, dependencies or user modifications, as they run as containerized Docker or Apptainer containers. This enables transparency and ensures computational reproducibility^10^. Two PET-BIDS Apps have been developed that perform preprocessing of the PET data: PETPrep^11^ and Pipeline Initializing Container for Neuro-Imaging Computations (PICNIC)^12^.

The radioligand [^11^C]UCB-J has seen a rapid increase in scientific interest since its development and validation^13,14^. This radioligand binds to the canonical synaptic vesicle glycoprotein 2A (SV2A) and the binding (as quantified as total volume of distribution (*V*_T_) or Binding Potential (*BP*_ND_)) is interpreted as a measure of synaptic density^14^. Changes in synaptic density as measured by [^11^C]UCB-J binding have been observed in schizophrenia^15^, multiple sclerosis^16^, Parkinson’s disease and dementia with Lewy bodies^17^. Synaptic density was also shown to be correlated with depression severity in a study by Holmes et al. (2019)^18^. Previous research has reported good test-retest performance in patients with Alzheimer’s disease^19,20^, and healthy controls^19–21^ however no studies have yet evaluated test-retest performance in subjects with psychiatric disorders.

Our primary objective with the current study was to assess the real-life performance of these recently developed PET-BIDS Apps on a dataset of [^11^C]UCB-J data collected from subjects with Major Depressive Disorder (MDD) in comparison to an in-house PET-processing software that has been optimized for the analysis of data collected at Karolinska Institutet, called Solena. *V*_T_ was chosen as the parameter of interest. Blood modeling and kinetic modeling was performed with the same open-source tool for all pipelines. Our secondary aim was to quantify test–retest statistics of [^11^C]UCB-J in this sample of subjects with MDD, who received niacin as a placebo control in a randomized clinical trial. Lastly, we aimed to investigate whether the choice of different preprocessing pipelines meaningfully influenced test-retest consistency.

## Material and Methods

### Study design and subjects

The study protocol was approved by the Swedish Ethical Review Authority (etikprö vningsmyndigheten) and the Swedish Medical Products Agency (lä kemedelsverket) under Dnr 2020-03700, and EudraCT-nr 2020-002790-94. The study protocol and clinical results have previously been published by Yngwe et al. (2026)^22^. Briefly, 35 subjects with recurrent MDD were randomized 1:1 in a blocked design to receive either a 25 mg dose of psilocybin or 100 mg of niacin as an active placebo, both with psychotherapeutic support. The support consisted of a preparation session the day before dosing, and integration sessions on day 1, 8 and 15 after dosing. Throughout the study, the Montgomery- Å sberg Depression Rating Scale (MADRS) was used to determine the severity of the depression^23^. All subjects provided written informed consent and the study was conducted in compliance to the Declaration of Helsinki. Only the data from the 18 subjects receiving niacin were considered for this study, and the final analysis included 14 subjects after the exclusions described below.

### [^11^C]UCB-J synthesis

[^11^C]UCB-J was prepared by reaction of the precursor, (R)-3-(difluoroboranyl)-4-((2-oxo-4-(3,4,5-trifluorophenyl) pyrrolidin-1-yl)methyl)-pyridin-1-ium fluoride, with carbon-11 labelled methyl iodide (MeI) via Pd-catalyzed carboncarbon bond formation using TracerMaker automated module (ScanSys Laboratorieteknik, Denmark), largely as described previously by Nabulsi et al. (2016)^13^. Further details on how the synthesis protocol deviates from Nabulsi et al. (2016) can be found in the supplementary materials.

### PET data acquisition

Subjects underwent two PET measurements with the radioligand [^11^C]UCB-J, one 1–7 days before dosing and the second 15 days (± 7) after dosing. All PET measurements were acquired between January 2021 and February 2023. One subject withdrew after dosing without completing the imaging protocol, resulting in 17 subjects with two PET examinations. While the study was not set up as a test-retest study, we used these repeated measurements to assess the reliability and consistency of *V*_T_.

A bolus injection was administered with a target dose of 400MBq/70kg and dynamic PET data were acquired for 90 minutes in list mode on the ECAT HRRT (High Resolution Research Tomograph, Siemens Molecular Imaging). A 6-minute transmission scan (using 137-Cs sources) was used for attenuation correction. Dynamic data were reconstructed into 37 frames using three-dimensional ordinary Poisson ordered subset expectation maximization (OP-3D-OSEM), including modeling of the system’s point spread function. The resulting frame durations were 8×15s, 8×30s, 4×60s, 5×120s, 10×300s, 2×600s. After reconstruction, PET measurements from 3 subjects with high motion (maximum displacement relative to the reference frame *>* 4mm as determined by SPM12 motion correction as implemented in Solena) were referred for additional re-reconstruction with coregistration to the attenuation scan for the primary planned PET analysis for comparison of psilocybin and niacin treated subjects. To avoid introducing additional variability from motion and re-reconstruction, only the 14 subjects who had acceptable motion (*<* 4mm) for both PET measurements were included.

### Arterial blood sampling

Arterial blood was sampled throughout the PET measurement from an arterial cannula inserted in the radial artery. During the first 10 minutes, an Automatic Blood Sampling System (ABSS; Allogg AB, Sweden) was used, and additional discrete manual samples were drawn for whole blood and plasma radioactivity, as well as metabolite analysis throughout the duration of the PET measurement. The parent fraction of [^11^C]UCB-J and its radiometabolites in plasma were measured using a protein precipitation procedure followed by reversed-phase radio-high-performance liquid chromatography analysis. Blood samples for metabolite analysis were collected at 2, 10, 20, 30, 45, 60, 75, and 90 minutes post-injection of [^11^C]UCB-J. Samples for whole blood and plasma radioactivity measurements were taken at 1, 2, 3, 5, 7, 10, 15, 20, 25, 30, 45, 60, 75 and 90 min. Details on metabolite analysis can be found in the supplementary materials.

### MRI data acquisition

T1-weighted (T1w) and Fluid-Attenuated Inversion Recovery (FLAIR) Magnetic Resonance Imaging (MRI) scans were collected on two different 3 Tesla scanners. The first subset of ten subjects underwent scanning on the General Electric (GE) Discovery MR750. The second subset of four subjects underwent scanning on the GE SIGNA Premier. Details about the acquisition parameters can be found in table S1. MRI data were acquired 7–35 days before and the day after treatment. Quality of the MRI scan was investigated using manual inspection of the T1w image. The first (pre-dosing) scan was used if deemed of acceptable quality via visual inspection. For one subject, substantial artifacts were present, and the second MRI scan was used instead.

### BIDS conversion

PET data stored in ECAT7 format were converted to NiFTI using PET2BIDS (pypet2bids) version 1.3.9^24^. A custom python script was created to update json files created by PET2BIDS with additional information not included in the PET header information. MRI data were converted from DICOM to NiFTI using dcm2bids version 3.1.1^44^, which uses dcm2niix version v1.0.20240202^25^. Blood data were converted to BIDS-compliant tsv files using a custom MATLAB script. The resulting dataset was verified as BIDS-valid using the command-line version of the bids-validator version 2.1.1^26^.

### Preprocessing

PET data were preprocessed using three different pipelines: an in-house software (Solena), as well as two open-source publicly available BIDS Apps (PETPrep and PICNIC). All three preprocessing pipelines included frame-to-frame motion correction of PET data, segmentation of the MRI data, coregistration and extraction of tissue Time Activity Curves (TACs) from each of the regions of interest. No correction for partial volume effects was applied. The most important differences between pipelines can be found in table 1. A longer description of the preprocessing for each tool can be found in the Supplementary methods.

**Table 1.** Difference between the different preprocessing pipelines.

|  | Solena | PICNIC | PETPrep |
|---|---|---|---|
| **General Implementation** |  |  |  |
| User interface | Graphical user interface | Docker container + input deck | Docker container |
| Implementation language | MATLAB | Python | Python |
| **MRI** |  |  |  |
| Sequence used | T1w | T1w | T1w (+ FLAIR) |
| Freesurfer version | 6.0.0 | 7.3.2 | 7.4.1 |
| Freesurfer extra options | “-brainstem-structures” segmentation<sup>27</sup> |  | “gtmseg” segmentation from PETSurfer<sup>28,29</sup><br>Output further refined using the ANTs brain segmentation using Mindboggle |
| **PET** |  |  |  |
| Motion correction | SPM12 realignment to the first 4 min | McFLIRT in FSL, registering to frame 8 (105–120s) | FreeSurfer’s mri robust register and tools from FSL, estimating motion from frame 9 (after 120s) and registering to an unbiased mean volume |
| Coregistration | SPM12 coreg with normalized mutual information to cropped MR | Lowest cost function based on FLIRT in FSL | Trimming using FSL robustfov, brainmasking, FreeSurfer’s mri coreg |
| Extraction of ROI volume | PET-space | MRI-space | MRI-space |

Since FLAIR MRI was not available for all subjects, we ran PETPrep both with and without the inclusion of the FLAIR. Our primary analysis was conducted using only the T1w MRI scans. However, we conducted a secondary analysis to compare these results with those generated by PETPrep when run with the inclusion of the FLAIR scan.

### Blood processing

Blood data were processed using the R packageBloodstream (version 0.1.0) which is based on the kinfitr R package^30^. The parent fraction and whole blood to plasma ratio were fit using hierarchical generalized additive models using a beta distribution and a student-*t* distribution, respectively, for their likelihood functions. No additional modeling was applied to the whole blood or plasma arterial input function, relying instead on piecewise linear interpolation.

### Kinetic modeling

Resulting TACs and arterial input functions were loaded into R version 4.4.3 with RStudio version 2024.12.1.563^31,32^. Kinfitr version 0.8.0 was used for kinetic modeling^30^. First, 9 volume-weighted composite regions of interest (ROIs) were created using the subregions listed in table S2, which were based on the ROI definition in Solena. Next, a composite region consisting of a volume-weighted average of all 9 ROIs was created and used to fit the delay between the arterial input function and the PET data. For this delay fit, based on a 1-Tissue Compartment Model (1TCM), the first 3 minutes of data were used, with each curve fitted 10 times with randomly selected starting points and a fixed blood volume of 5% and with uniform frame weights. Frame weights for subsequent steps were calculated using the same composite ROI. Next, a 1TCM was fit to all ROIs using the delay estimate, weights, as well as the arterial input function and whole blood curves created by bloodstream, using a fixed blood volume of 5%. Each curve was fitted 20 times with randomly selected start parameters to reduce the likelihood of estimating local minima.

### Quality control

The resulting html quality control reports generated by PETPrep, PICNIC and Bloodstream were checked manually by GH. For preprocessing in Solena, all quality control was carried out using visual inspection during preprocessing by MA. While a small segmentation defect was present for one subject for PICNIC, this was considered small enough that no manual intervention was needed for any pipeline.

### Statistics

Results from the different pipelines (extracted ROI volumes and *V*_T_) were compared using Pearson’s correlation coefficients as well as linear regression, using the lm function from the stats package in R version 4.4.3. Since slight differences in volumes between PET sessions were possible for Solena, due to extraction of the ROI volumes in PET space, we took an average volume across sessions for comparison with the other tools. The other tools provided only one ROI volume for both sessions. For *V*_T_, correlations were not corrected for repeated measures, and the 28 measurements were instead entered into the model as independent data points. We calculated the two-way mixed effects single rater Intraclass Correlation Coefficient (ICC) for consistency (ICC_c_) of *V*_T_ estimated between pipelines^33^, using the irr package^34^ in R. ICC_c_ values were calculated pairwise for the three tools. Different hierarchical parent fraction models (with slopes for (PET) or (PET + subject) or (PET + subject + session)) were compared using the Akaike information criterion as implemented in the stats package, with the lowest value indicating the preferred model. In these models “PET” refers to each individual scan (N=28).

To assess the within-individual consistency of estimates from the different preprocessing pipelines, we calculated test-retest statistics for each of the different tools. The difference between session 1 and session 2, expressed as the Absolute Percentage Difference (APD) and the signed Percentage Difference (PD) was calculated using equations 1 and 2, with the test measurement being session 1 and the retest measurement being session 2. For comparisons between tools, the PD is referred to as bias. We also calculated the two-way mixed effects single rater ICC with absolute agreement (ICC_a_) for test-retest reliability^33^ using the irr package^34^ in R. Systematic differences in *V*_T_ between PET imaging sessions for each of the pipelines were assessed using a two-tailed, paired *t*-test. From the model, we also extracted the proportional standard error (standard error expressed as a proportion of the estimate) of *V*_T_ as an estimate of model fitting accuracy, and compared differences between sessions for each tool with a two-tailed paired *t-* test. We also calculated the area under the curve (AUC) for both the TACs and the arterial input function using the DescTools package, and corrected this for the injected dose (injected radioactivity/kg) to obtain the AUC-SUV. For the arterial input function, the curves were cropped to 75 minutes, as this was the latest timepoint where radioactivity measurements were available for all measurements. Differences in AUC-SUV for both TACs and arterial input function between sessions were assessed using a two-tailed, paired *t*-test. For all analyses, p-values *<* 0.05 were deemed significant. Due to the exploratory nature of this part of the analysis, we did not perform correction for multiple comparisons.

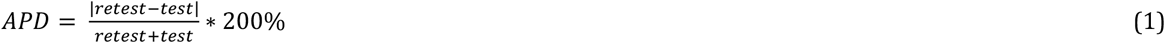

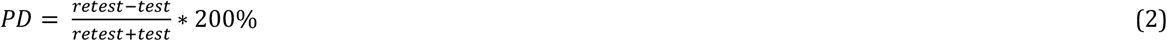

As our primary ROIs for comparison across tools we chose the frontal cortex, striatum, and hippocampus because of their importance in psychiatric research and their differences in anatomical localisation and shape. Lower binding of [^11^C]UCB-J in dorsolateral prefrontal cortex and hippocampus has previously been shown to be linked to depression severity^18^, and [^11^C]UCB-J binding in hippocampus has been linked to schizophrenia symptom severity^35^. Since the cortex and subcortical structures might have different susceptibilities in variation arising from differences in preprocessing, we included both the frontal cortex as a cortical example, and two subcortical structures with different shapes.

To assess the extent to which inclusion of the FLAIR scan could improve performance, we also compared outputs generated by PETPrep run with and without FLAIR scans. We performed two-tailed paired *t*-tests on the subset of subjects that had a FLAIR scan in the raw dataset (N=9). Comparisons for volumes included only the first sessions (N=9), while for *V*_T_ estimates both PET measurements were used (N=18), without correcting for repeated measures from the same subject. We considered *p*-values *<* 0.05 as significant, and no correction for multiple comparisons was applied.

## Results

The 14 subjects included in the analyses had a mean age of 41.5 (range 22–61), and included 10 females and 4 males. MADRS did not differ between session 1 and session 2, with a summed score of 25.4 ± 4.4 and 25.3 ± 5.5 (p=0.96), respectively.

The 28 measurements from the 14 subjects included in this analysis had a between measurement interval of 13 to 28 days (17.9±6.0, mean±sd). Injected radioactivity and injected dose did not differ significantly between pre- and post-treatment sessions, with an average injected radioactivity of 397 ± 68.9 MBq and 405 ± 75.8 MBq (p = 0.341) and injected dose of 5.50 ± 0.94 MBq/kg and 5.83 ± 0.18 MBq/kg (p = 0.189), respectively. However, a difference in injected mass between pre- and post-treatment sessions was observed, with an average injected mass of 0,86 ± 0,38 µg and 0,63 ±0,19 µg respectively (p = 0.028). Hence, a difference in specific activity was also observed between pre- and post-treatment sessions, with a specific activity of 548 ± 238 MBq/µg and 690 ± 184 MBq/µg respectively (p = 0.020).

### Quality control

The quality control reports from the PET pre-processing showed accurate coregistration with the MRI scan and no issues with motion correction. For PICNIC, a segmentation error for one subject in the delineation of the cerebellar cortex was identified, where the defined region included surrounding tissue. There were no issues with the modelling of the parent fraction or blood plasma ratio.

### Extracted ROI volumes

Since volumes from Solena were extracted in PET-space, and correlation between session 1 and session 2 was very high (all R^2^ > 0.98), we used the average volume between sessions for Solena. Correlation of volumes between the three preprocessing pipelines was high on average, with the lowest correlation of volume and high bias found for brainstem, see table 2. Direct comparison of volumes between tools can be found in figure S1-S3, and the direct comparison between sessions for Solena in figure S4.

**Table 2.** Bias and correlation coefficients (R^2^) for volume of the regions of interest between tools. An average volume between sessions was calculated for Solena, and all comparisons include all subjects (N=14). Bias estimates are calculated so that a positive bias corresponds to a higher value in the tool/session that is mentioned second. Comparisons with an R^2^ < 0.9 in bold.

|  | Solena ses-01 vs. ses-02 |  | Solena vs. PICNIC |  | Solena vs. PETPrep |  | PETPrep vs PICNIC |  |
|---|---|---|---|---|---|---|---|---|
| Region | Bias | R<sup>2</sup> | Bias | R<sup>2</sup> | Bias | R<sup>2</sup> | Bias | R<sup>2</sup> |
| Frontal cortex | 0.2% | > 0.99 | 0.2% | 0.99 | -4.1% | 0.95 | 4.3% | 0.98 |
| Parietal cortex | 0.0% | > 0.99 | -0.1% | > 0.99 | -4.6% | 0.98 | 4.4% | 0.98 |
| Temporal cortex | 0.0% | > 0.99 | 0.3% | > 0.99 | -2.2% | 0.96 | 2.5% | 0.98 |
| Occipital cortex | -0.1% | > 0.99 | 2.0% | 0.97 | -5.9% | 0.97 | 7.9% | 0.95 |
| Striatum | 0.2% | > 0.99 | 0.4% | 0.96 | -1.1% | 0.98 | 1.6% | 0.97 |
| Hippocampus | 0.5% | 0.98 | 1.3% | 0.92 | 5.1% | 0.92 | -3.8% | 0.95 |
| Thalamus | 0.0% | > 0.99 | 2.0% | 0.95 | 2.5% | 0.97 | -0.5% | 0.97 |
| Cerebellar cortex | -0.1% | > 0.99 | -1.0% | 0.96 | -2.0% | 0.98 | 1.0% | 0.98 |
| Brainstem | 0.0% | > 0.99 | -1.6% | 0.96 | **-54.5%** | **0.86** | **53.1%** | **0.90** |

### Direct TAC comparison

Differences in TAC curve values were assessed directly by calculating the percentage difference between tools, see figure 1. Because the first frame included very low and zero values, differences were only assessed from the second frame onward. Differences between tools varied widely between regions, but no clear pattern across time frames and regions were identified.

**Figure 1.**
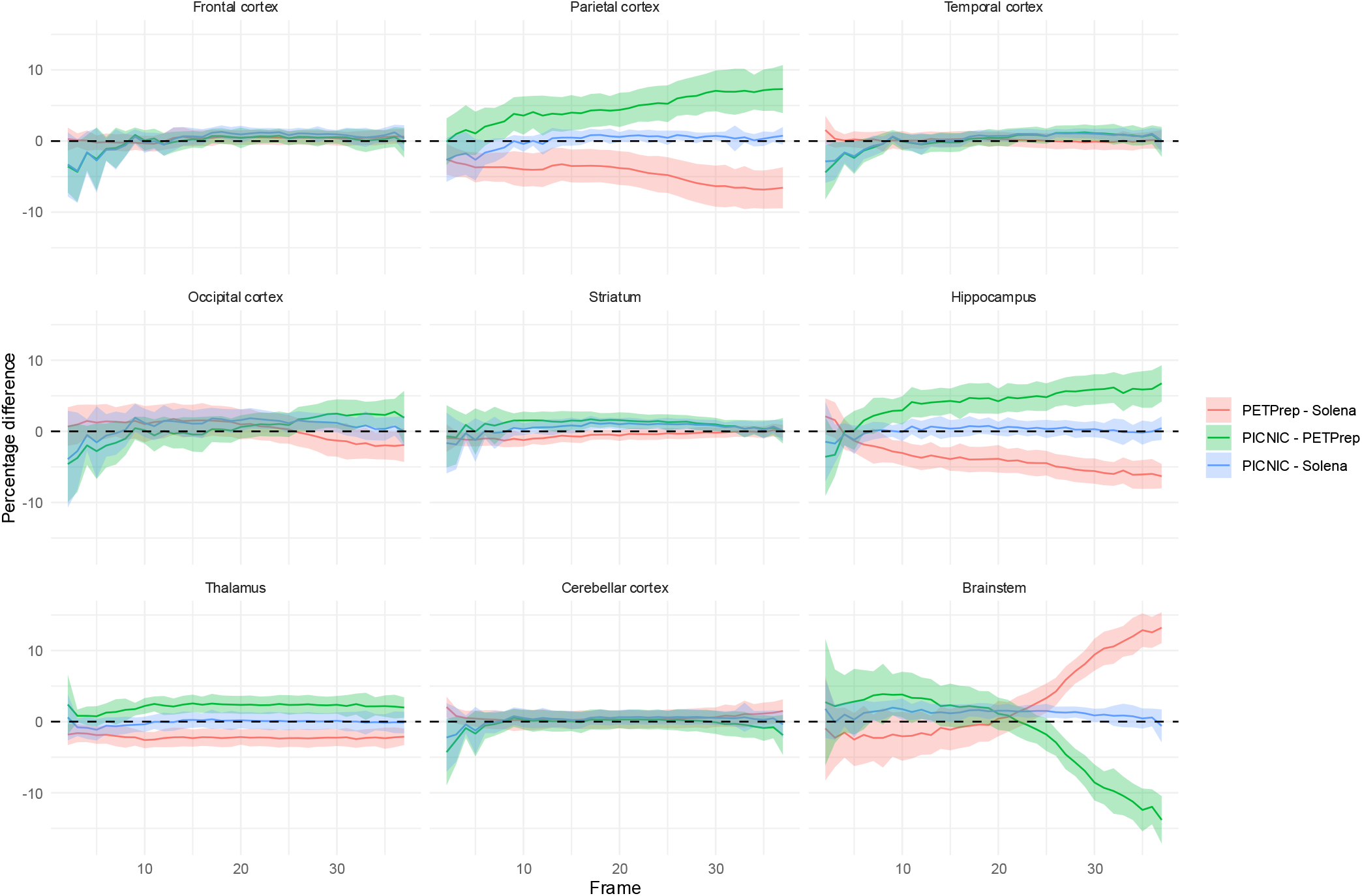
The percentage difference across regions and frames 2-37 for the three tool comparisons. With mean in bold and shaded areas representing one standard deviation around the mean

### Volumes of distribution

*V*_T_ showed generally high correlation between tools for all regions. In figure 2 results for three representative ROIs are reported. Correlations and ICC_c_ were very high between Solena and PICNIC, with all R^2^ and ICC_c_ estimates being ≥0.98, see table 3. While both R^2^ and ICC_c_ were slightly lower for PETPrep compared both to Solena and PICNIC, most R^2^ and ICC_c_ were still *>*0.95, with only a few exceptions. The lowest correlation was observed for parietal cortex for PETPrep when correlated with either Solena or PICNIC (R^2^ = 0.89 and 0.88, respectively). A comparison between tools for all regions is reported in figures S5-7.

**Table 3.** Bias, correlation coefficients (R^2^) and ICC_c_ (95% CI) for V_*T*_ as derived by the three different tools Solena, PICNIC and PETPrep. N=28 for all comparisons. Bias estimates are calculated so that a positive bias corresponds to a higher value in the latter tool. Comparisons with an R^2^ *<* 0.95 in bold.

|  | Solena vs. PICNIC |  |  | Solena vs. PETPrep |  |  | PETPrep vs. PICNIC |  |  |
|---|---|---|---|---|---|---|---|---|---|
| Region | Bias | *R*<sub>2</sub> | ICC<sub>c</sub> | Bias | *R*<sub>2</sub> | ICC<sub>c</sub> | Bias | *R*<sub>2</sub> | ICC<sub>c</sub> |
| Frontal cortex | 1.4% | 0.99 | 0.99 (0.99 - 1.00) | 0.7% | 0.99 | 0.99 (0.99 - 1.00) | 0.6% | 0.98 | 0.99 (0.98 - 1.00) |
| Parietal cortex | 1.0% | >0.99 | 1.00 (0.99 - 1.00) | -7.9% | **0.89** | 0.94 (0.88 - 0.97) | 8.9% | **0.88** | 0.94 (0.86 - 0.97) |
| Temporal cortex | 1.6% | >0.99 | 1.00 (0.99 - 1.00) | -0.2% | 0.98 | 0.99 (0.98 - 1.00) | 1.8% | 0.98 | 0.99 (0.98 - 1.00) |
| Occipital cortex | 0.8% | 0.99 | 0.99 (0.98 - 1.00) | -3.1% | **0.94** | 0.97 (0.93 - 0.99) | 3.9% | 0.95 | 0.97 (0.95 - 0.99) |
| Striatum | 0.8% | >0.99 | 1.00 (0.99 - 1.00) | 0.4% | >0.99 | 1.00 (0.99 - 1.00) | 0.4% | >0.99 | 1.00 (0.99 - 1.00) |
| Hippocampus | 0.1% | 0.98 | 0.99 (0.98 - 1.00) | -7.5% | 0.97 | 0.98 (0.96 - 0.99) | 7.6% | **0.95** | 0.97 (0.94 - 0.99) |
| Thalamus | 0.1% | 0.98 | 0.99 (0.98 - 1.00) | -2.3% | 0.99 | 0.99 (0.98 - 1.00) | 2.4% | 0.98 | 0.99 (0.98 - 1.00) |
| Cerebellar cortex | 0.6% | >0.99 | 1.00 (0.99 - 1.00) | 0.9% | 0.99 | 0.99 (0.98 - 1.00) | -0.3% | 0.99 | 0.99 (0.98 - 1.00) |
| Brainstem | 1.1% | >0.99 | 1.00 (0.99 - 1.00) | 10.2% | 0.97 | 0.98 (0.95 - 0.99) | -9.1% | 0.96 | 0.98 (0.95 - 0.99) |

**Figure 2.**
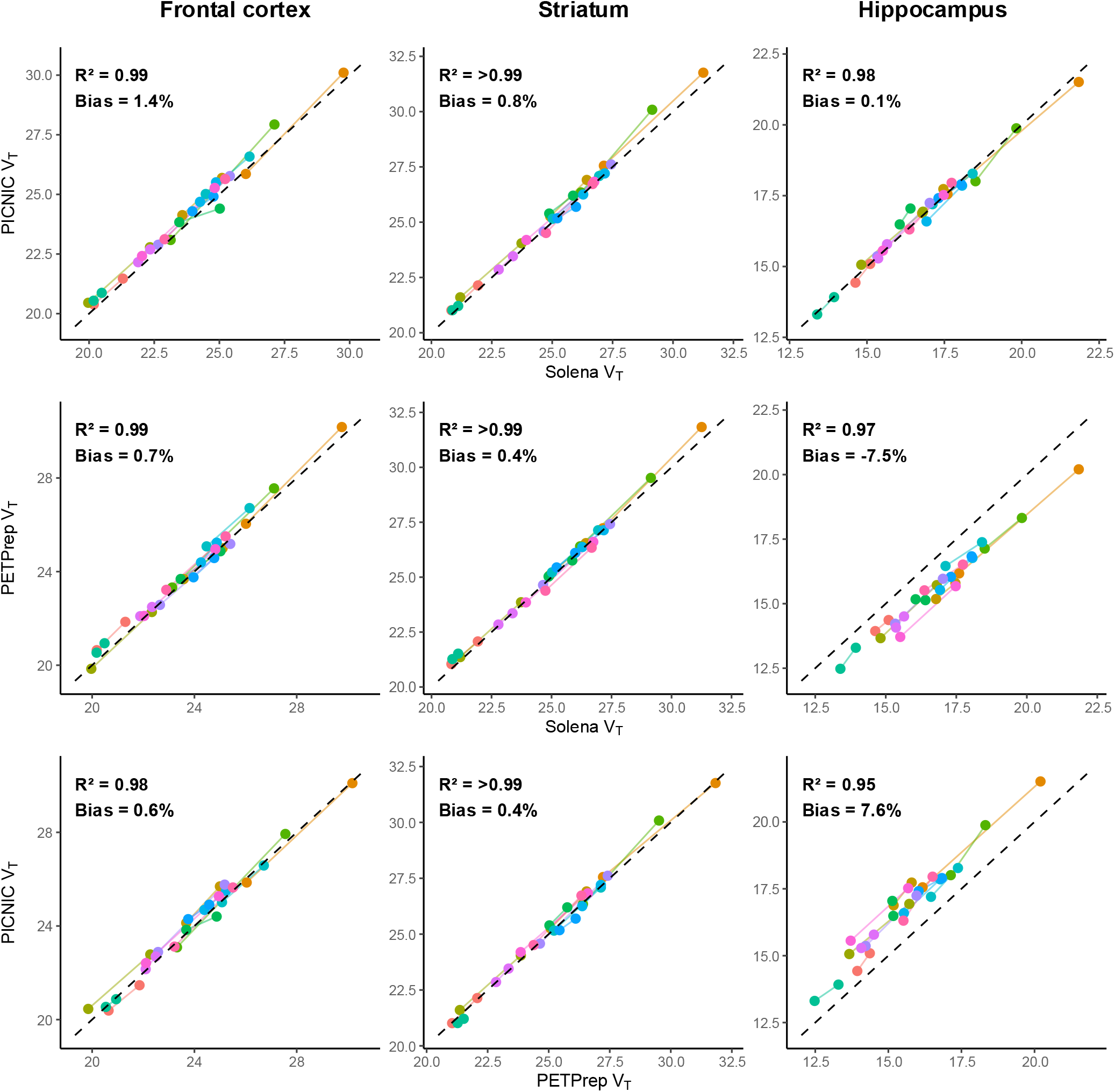
Direct comparison of *V*_T_ as derived by the three different preprocessing tools for frontal cortex, hippocampus and striatum. Top row: comparison between Solena and PICNIC. Middle row: comparison between Solena and PETPrep. Bottom row: comparison between PETPrep and PICNIC. Bias is calculated using the PD formula, with positive bias indicating higher *V*_T_ for the tool on the y-axis relative to the tool on the x-axis. Data points are color-coded by subject, colored lines connect the two measurements of the same subject.

To investigate the influence of the inclusion of FLAIR data on results generated by PETPrep, we reran PETPrep as described previously without the “--ignore flair” tag. This automatically includes any FLAIR scans found in the BIDS dataset to provide additional information for delineation of ROIs using FreeSurfer. Resulting ROIs volumes and *V*_T_ values are reported in figure S8. Inclusion of FLAIR led to smaller volumes for regions in the cerebral cortex, and slightly larger volume for cerebellar cortex, see figure S8A. *V*_T_ was significantly increased in all regions except brainstem (uncorrected p-values *<*0.05, see table S3 and figure S8B); however, the magnitude of these differences was small with an average *V*_T_ increase of 0.5% (range 0.1–1.0%) across regions.

#### Test-retest statistics

When comparing the *V*_T_ values for the test (session 1) and retest (session 2) measurements, lower mean values for all regions were found for the retest measurement, across the three tools. Similar average PD and APD values across regions were observed for PETPrep (PD: ™4.67%, APD: 7.33%), Solena (PD: ™4.73 APD: 7.43%), and PICNIC (PD: ™4.88% APD: 7.56%), respectively. Values for three representative regions for the three tools are reported in table 4. All regional values for the tools separately can be found figure S9 and in tables S4-6. As reported in table 4, ICC_a_ for two-way mixed effects and absolute agreement between sessions and PD/APD estimates are similar across tools.

**Table 4.**
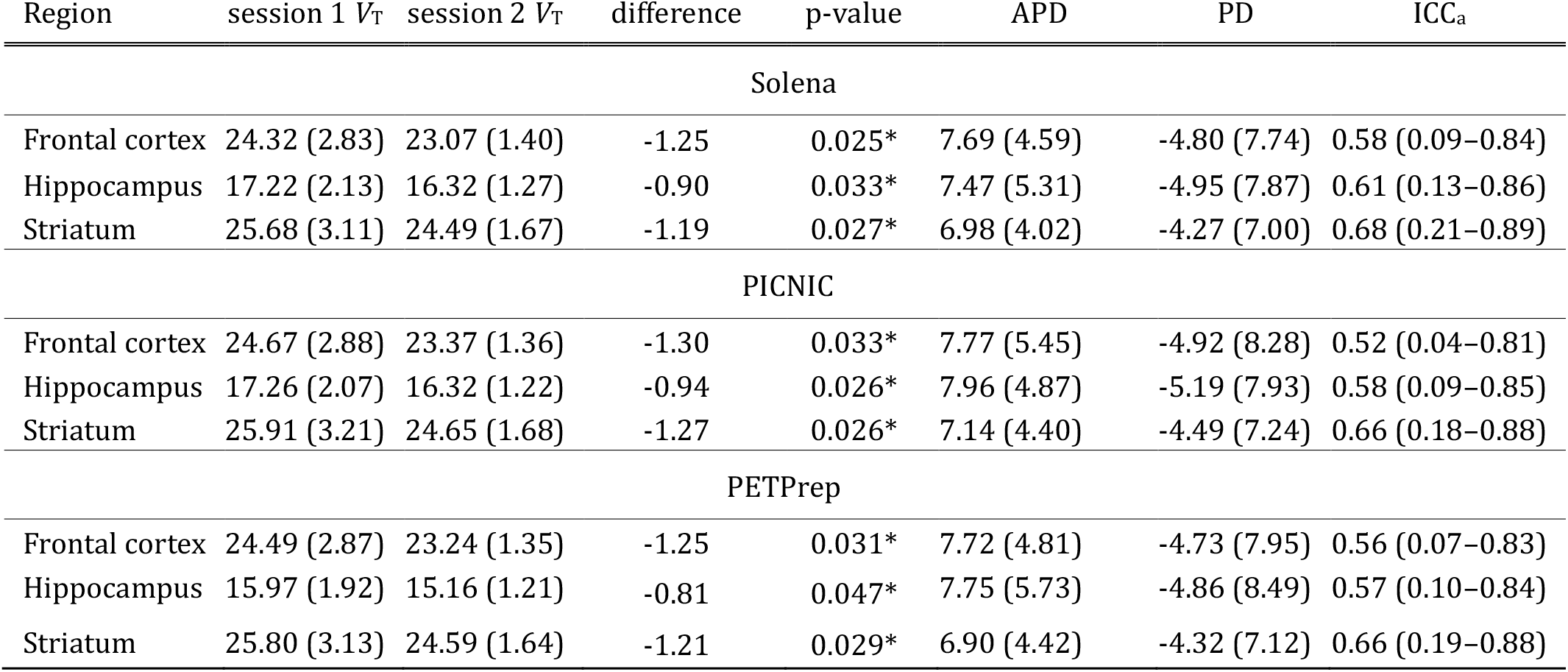
Comparison of *V*_T_ as derived using Solena, PICNIC and PETPrep for three representative ROIs for session 1 and session 2. Comparison between sessions is represented as difference (session 2 - session 1), APD and PD, reported as mean (sd). ICC_a_ calculated with two-way mixed effects and absolute agreement and reported as estimate (95% CI). P-values are calculated using a paired two-tailed t-test. P-values *<* 0.05 are indicated with *.

| Region | session 1 *V*<sub>T</sub> | session 2 *V*<sub>T</sub> | difference | p-value | APD | PD | ICC<sub>a</sub> |
|---|---|---|---|---|---|---|---|
| Solena |  |  |  |  |  |  |  |
| Frontal cortex | 24.32 (2.83) | 23.07 (1.40) | -1.25 | 0.025* | 7.69 (4.59) | -4.80 (7.74) | 0.58 (0.09–0.84) |
| Hippocampus | 17.22 (2.13) | 16.32 (1.27) | -0.90 | 0.033* | 7.47 (5.31) | -4.95 (7.87) | 0.61 (0.13–0.86) |
| Striatum | 25.68 (3.11) | 24.49 (1.67) | -1.19 | 0.027* | 6.98 (4.02) | -4.27 (7.00) | 0.68 (0.21–0.89) |
| PICNIC |  |  |  |  |  |  |  |
| Frontal cortex | 24.67 (2.88) | 23.37 (1.36) | -1.30 | 0.033* | 7.77 (5.45) | -4.92 (8.28) | 0.52 (0.04–0.81) |
| Hippocampus | 17.26 (2.07) | 16.32 (1.22) | -0.94 | 0.026* | 7.96 (4.87) | -5.19 (7.93) | 0.58 (0.09–0.85) |
| Striatum | 25.91 (3.21) | 24.65 (1.68) | -1.27 | 0.026* | 7.14 (4.40) | -4.49 (7.24) | 0.66 (0.18–0.88) |
| PETPrep |  |  |  |  |  |  |  |
| Frontal cortex | 24.49 (2.87) | 23.24 (1.35) | -1.25 | 0.031* | 7.72 (4.81) | -4.73 (7.95) | 0.56 (0.07–0.83) |
| Hippocampus | 15.97 (1.92) | 15.16 (1.21) | -0.81 | 0.047* | 7.75 (5.73) | -4.86 (8.49) | 0.57 (0.10–0.84) |
| Striatum | 25.80 (3.13) | 24.59 (1.64) | -1.21 | 0.029* | 6.90 (4.42) | -4.32 (7.12) | 0.66 (0.19–0.88) |

Next, we investigated if the negative test-retest bias was associated with differences in the metabolism (parent fraction) or uncertainty of the estimated *V*_T_ from the one tissue compartment model. Visually, there was only a small difference in the parent fraction between the two sessions as can be seen in figure 3, with slightly faster metabolism in session 2. The Cohen’s dz showed an estimated moderate effect at later timepoints, though confidence intervals were wide, see figure S10. The Akaike information criterion showed that from models including “PET” or “subject and PET”, or “subject, session and PET”, using “subject and PET” was preferred, implying that systematic session differences did not contribute meaningfully to model performance for describing the metabolite parent fraction. There were no differences in the standard error of the estimated *V*_T_ between sessions for any region across any of the tools see table S7, indicating modelling uncertainty was not significantly different between sessions. We observed an increase in the AUC of the arterial input function in the second session (2452 vs. 2630, p-value 0.016), see figure S11. While AUC of the regional TACs were numerically higher in the seconds session, no significant differences were observed for any region in any tool, see table S8.

**Figure 3.**
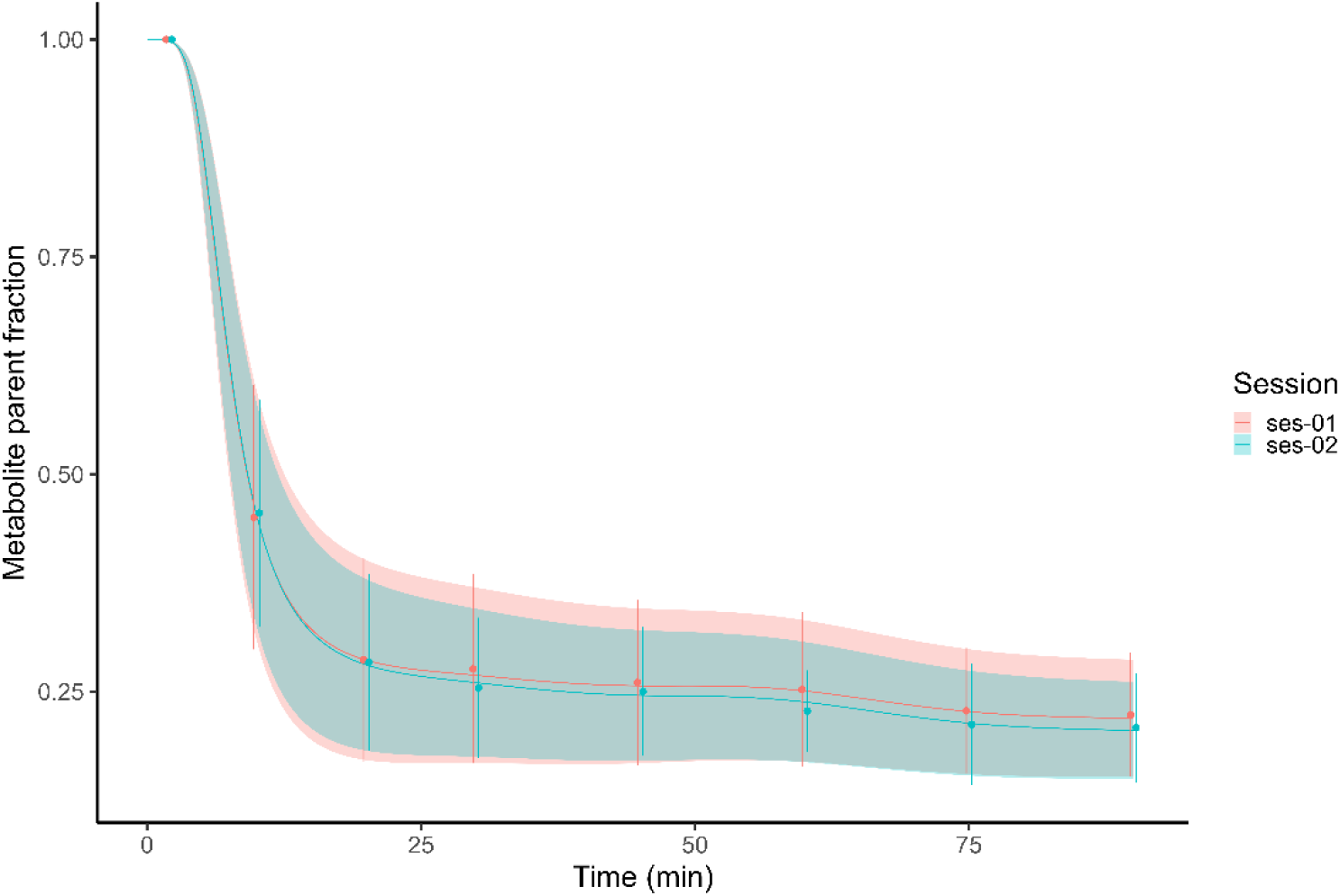
Measured and fitted values for the parent fraction per session. For session 1, one subjects data had to be extrapolated due to missing data at 90 minutes. Discrete time points include 11-14 measurements each. Positions of discrete points are dodged to allow for better visual separation, but occurred at the same time points for both sessions

## Discussion

In this study we validated the use of BIDS Apps for the preprocessing of PET image data for consistency between tools, as well as reproducibility and reliability within subjects across measurements. The three pipelines produced generally consistent outcomes with respect to both ROI volumes and *V*_T_ values. R^2^ values between tools for *V*_T_ were *>* 0.85, with most regions having a R^2^ *>* 0.95. ICC_c_ indicated excellent consistency for most tools and regions^33^, only showing good-excellent consistency for parietal cortex for PETPrep compared to the other tools, with estimated ICC_c_ (95% CI) of 0.94 (0.88–0.97) for Solena and 0.94 (0.86–0.97) for PICNIC, respectively.

The main differences in ROI volume and *V*_T_ estimates between tools are likely due to the differences in segmentation. Most importantly, PETPrep makes use of a different FreeSurfer segmentation (gtmseg) compared to the other tools (aparc+aseg). This gtmseg uses a higher resolution (0.5×0.5×0.5mm instead of 1×1×1mm) and includes a different segmentation of brainstem and cerebellum. In line with that, brainstem volumes were less consistent for PETPrep versus the other tools (*R*^2^ values of 0.86 and 0.90). The gtm segmentation in PETPrep excludes the pons from the brainstem region, while the other tools include the pons. Indeed, absolute brainstem volumes from PETPrep showed an average bias of -53.1% compared to those extracted by PICNIC (see table 2 and figure S3), corresponding to a mean decrease of 42% in volume. However, the inconsistency in brainstem volume between tools did not seem to affect the consistency of the estimation of *V*_T_, as the correlation with the other tools still yielded *R*^2^ values of 0.97 and 0.96 although a bias of ∼10% was observed. On the other hand, differences in segmentation of the cerebellum (which excludes vermis for PETPrep) did not lead to lower correlations or ICC_c_ in volume or *V*_T_ with the other tools. The region-dependant bias warrants caution for direct comparison of *V*_T_ between tools in future studies, and researchers are advised to use the same BIDS App for processing of all subjects within the same study to prevent the introduction of bias.

Additional variability between tools can be explained by other differences between the preprocessing pipelines. For example, the three tools use different versions of FreeSurfer, which is known to influence regional volumes^36,37^. This implies that, while the regions have the same name, the actual definition of these regions differs between tools. Another potential source of differences in volumes and *V*_T_ could be that PETPrep uses ANTS and FSL for skull stripping and brain extraction within the FreeSurfer pipeline, while the other tools make use of the default FreeSurfer procedures. While correlation coefficients for ROIs volumes and *V*_T_ values were still high across the regions included in this study, we cannot exclude the possibility that the differences in segmentation, Freesurfer version and other differences between pipelines could lead to differences in both volume and *V*_T_ when smaller or different subregions are analyzed.

We also investigated the effect of the inclusion of FLAIR in PETPrep, since PETPrep automatically detects the presence of FLAIR scans and uses this additional information to improve definition of the pial surface. Since FLAIR scans were only available for 9 subjects, this was performed as a secondary analysis. Inclusion of FLAIR gave rise to cortical regions with a lower volume, and cerebellar cortex volume increasing with FLAIR usage. The reductions in cortical volumes were expected, as the improvement of the pial surface should exclude voxels that include cerebrospinal fluid and other extracerebral structures. In turn, this led to significant increases in *V*_T_ in all regions except brainstem, but absolute differences were very small. These differences are close to negligible for most applications, as they are much smaller than the within-individual or between-individual variability. However, it is increasingly important to understand these small sources of bias, as tools like these enable harmonised analysis of multi-centre datasets with large sample sizes, for which bias becomes an increasingly important consideration.

Consistently across tools, we found lower estimates for [^11^C]UCB-J *V*_T_ in the retest measurement relative to the test measurement (see figure S9). A small difference in plasma parent fraction was present between sessions, without differences in fitting accuracy of the one tissue compartment model, and we deem it unlikely that the parent fraction resulted in the lower *V*_T_. In line with the session effect for *V*_T_, we observed an increase in arterial input AUC but not TAC AUC. There was also a difference between sessions in injected mass and specific activity, but since inject mass was lower in the second scan, and specific activity was higher, we deem it implausible that this would have resulted in a lower *V*_T_ A negative bias has been found in other studies in healthy controls^20^; however, we note that the current study was not primarily designed as a test-retest study. Other causes of variability, including some effect of niacin, the psychotherapeutic framework and placebo or nocebo effects on [^11^C]UCB-J binding, cannot be excluded. Since no differences in mean MADRS between the two PET measurements were present, we consider it unlikely that this change in [^11^C]UCB-J binding is related to biological changes due to depression severity. Within the study timeframe we do not see other plausible biological mechanisms that could cause a decrease in SV2A availability or density. While some evidence exists for an association between depression duration and hippocampal volume loss^38^, and synaptic loss that can be present in depression^39^ could have continued during the study duration, we think it is implausible that this would lead to a 5% decrease in synaptic density across only three weeks. The presence of a negative bias of [^11^C]UCB-J binding in retest measurements has been reported previously. For example, Tuncel et al. (2022) and Tuncel et al. (2021) have previously reported a negative bias of [^11^C]UCB-J binding with APD values in the same range as reported here^19,20^. However, Khattar et al. (2026) showed no negative bias in 30 healthy subjects undergoing same-day or longitudinal [^11^C]UCB-J test-retest measurements^40^. While we consider it possible that the negative bias observed in the current study is merely a spurious finding on account of our small sample size, pooling of [^11^C]UCB-J test-retest data across centres could provide a good opportunity to use BIDS Apps to evaluate this question in a more comprehensive and standardized manner.

Point estimates for ICC_a_ ranged from 0.52–0.71 across regions for PICNIC, between 0.56–0.7 for Solena, and 0.56– 0.72 for PETPrep. However the large confidence intervals indicates the ICC_a_ is consistent with anything from poor to good test-retest reliability^33^. Our estimates of the ICC_a_ values were numerically lower than those previously reported by Finnema et al. (2018)^21^, however differences in populations (healthy controls vs. MDD), sample size (5 vs. 14) and between measurement intervals (same day vs. 2–4 weeks) make direct comparisons difficult. Preprocessing pipelines had little to no influence on test-retest metrics, as both APD, PD and ICC_a_ were highly similar across tools.

The current study has several limitations. Firstly, it only included data from one PET camera and one radioligand, collected at one study site. We also only assessed *V*_T_ estimates, as we could not obtain a centrum semiovale from PICNIC which is the preferred reference region for [^11^C]UCB-J to estimate *BP*_ND_^14,40,41^. This limits the generalisability of these results, and we cannot say that the three pipelines will produce the same results when applied to data from other radioligands. As [^11^C]UCB-J distribution throughout the brain is likely well suited for automatic motion correction, it would be very interesting to see how well these tools perform for radioligands that have a less favorable distribution pattern for motion correction or coregistration with the MRI scan. However, it is encouraging that automated preprocessing by BIDS Apps performed similarly to an inhouse software optimized for our data collection procedures. Secondly, our sample size was relatively small, and it is possible that more issues with automated preprocessing are encountered in larger datasets from a less homogeneous sample of subjects. Thirdly, we used the BIDS Apps with default settings for PETPrep and using a PICNIC “input deck” similar to the example provided by PICNIC. While we deemed this to be the fairest comparison from a real-world usage perspective, more consistent results between tools could potentially be achieved by choosing alternatives for some of the preprocessing steps that are more similar across tools. However, testing the full range of possible settings was beyond the scope of the current investigation.

Both BIDS Apps produce comprehensive quality control reports which facilitate quality control procedures, and record all settings used for performing the analytical procedures in either the reports or in JSON files. This simplifies identification of issues which might otherwise have required specialised skills to evaluate. This also means that for radioligands with a less favourable distribution pattern, for which the default parameters might be less suited, researchers are able to identify these issues and to adapt and optimize analysis parameters where necessary. Similarly, manual intervention is also possible in cases where minor errors do persist. For example, one subject processed with PICNIC had a segmentation error of the cerebellar cortex, which led to the inclusion of part of the confluence of sinuses, extending into the superior sinus sagittalis in the cerebellar cortex ROI. This led to a slightly higher volume and lower estimated *V*_T_ for this subject compared with the other preprocessing pipelines (see figure S1 and S6, cerebellar cortex, dark blue). While manual intervention could have increased the consistency with the other tools, this also comes at the expense of full reproducibility.

Overall, we consider the results from this study encouraging. While our in-house software is optimized to preprocess data collected at our PET-center, the correlations and ICC_c_ with the evaluated BIDS Apps were high, and performance on test-retest metrics were very similar. Together, these results suggest that the general and non-catered design of BIDS Apps can yield robust performance for PET analysis. Reproducibility is a well-recognized challenge in neuroimaging, and automated, standardized preprocessing pipelines can help facilitate the generation of more comparable outcomes, especially from the perspectives of data sharing and multi-center collaboration. In cases where raw data cannot be shared due to local data protection regulations, running these BIDS Apps locally can ensure that preprocessing is kept consistent across sites, eliminating the risk of inconsistent local software environments or different user choices during preprocessing. Completion of the BIDS standard for PET derivatives (BIDS extension BEP023) will make the integration of outputs from different tools even more seamless, which will also simplify the application of BIDS Apps for kinetic modeling, such as PETFit^42^ and PETSurfer^43^, to further automate and standardize the analysis of PET data.

## Conclusion

PET BIDS Apps can reliably and reproducibly extract TACs from raw PET image data stored according to BIDS. Usage of these TACs for kinetic modeling leads to estimations of *V*_T_ which are both consistent with one another, as well as consistent with our in-house software which is specifically tailored for use with our in-house data. Bias between tools was present and differed across regions. Quantification of test-retest variability was highly similar across tools, indicating a similar sensitivity to intra- and inter-subject variability. Wider implementation of PET BIDS Apps can improve comparability and reproducibility across studies.

## Supporting information

Supplement_to_BIDS_apps_validation_preprocessing_of_UCBJ_data

## Acknowledgments

We thank the members of the PET-research group at Karolinska Institutet for their valuable discussions and suggestions on this manuscript. We thank Aleksander Lebedev, Kasim Acar and Predrag Petrovic at Karolinska Institutet for their contributions to collecting and sharing MRI-data used in this project. We also thank Eric Hauser for his contributions to the PICNIC software.

## Author Contributions

## Disclosure/conflict of interest

### Ethical Considerations

This study was approved by the Swedish Ethical Review Authority (etikprö vningsmyndigheten) and the Swedish Medical Products Agency (lä kemedelsverket) under Dnr 2020-03700, and EudraCT-nr 2020-002790-94 on September 16th, 2020.

### Consent to participate

All participants provided written informed consent.

### Consent for publication

Not applicable.

### Declaration of conflicting interest

The author(s) declared no potential conflicts of interest with respect to the research, authorship, and/or publication of this article

### Funding

Osmond Labs provided practical support. The foundation Norrsken Mind provided funding. The Swedish Research Council provided funding (grant 2021-01750). Granville Matheson was supported by the Swedish Research Council (2020-06356 and 2024-03534) and Strategic Research Area Neuroscience (StratNeuro). Mikael Tiger was supported by Region Stockholm (clinical research appointment). Usona Institute provided the investigational product, the treatment manual and a web-based training.

### Data availability

The data that support the findings of this study are available on request from the corresponding author and subject to approval by the relevant ethics committee/institution. The data are not publicly available due to privacy or ethical restrictions.

## Notes

### Competing Interest Statement

The authors have declared no competing interest.

