## Supplement_to_BIDS_apps_validation_preprocessing_of_UCBJ_data for "A Comparison and Validation of BIDS Apps for Preprocessing of [^11^C]UCB-J PET data"

### Supplementary methods and results

#### *Supplementary methods*

*[11C]UCB-J synthesis* In brief, 0.5–0.8 mg of tris(dibenzylideneacetone)dipalladium(0) (Sigma-Aldrich, USA), 0.5–0.8 mg of tri-1-napthylphosphine (Sigma-Aldrich, USA), and 2.4–3.1 mg potassium carbonate (Sigma-Aldrich, USA) were mixed with 500 µl of DMF/water solution (8:1) and [$^{11}$C]methyl iodide obtained from reaction of cyclotron-produced [$^{11}$C]methane with iodine bubbled through the reaction mixture at +5 ° C. To the resulting solution ca. 0.5–0.7 mg of pre-hydrolysed precursor (PharmaSynth AS, Estonia) dissolved in 350–400 µl of DMF/water (8:1) was added and the mixture heated to 90° C for 5 min.

Precursor was hydrolysed to form free boronic acid intermediate by dissolving the amount weighted in 80 µl of 0.35 M HCl in MeOH, flushing the vial with argon and letting it stand at RT for 20-30 min, after which methanol/HCl mixture was removed under reduced pressure with Ar flow to dryness, and the residue redissolved in 350–400 µl of DMF/water solution (8:1), which was then used in the labelling procedure.

Reaction mixture was then cooled to ca. 40° C, and diluted with 2.5 ml of 2 M HCl 1:9 MeCN/water solution. The crude reaction mixture containing [$^{11}$C]UCB-J was purified using ACE 5 C18-HL HPLC column (ACE Chromatography) and MeCN/Ammonium formate 0.1 M pH 9 as the mobile phase. [$^{11}$C]UCB-J fraction was collected, diluted in 50 ml of water, and [$^{11}$C]UCB-J was then isolated using solid-phase extraction employing Waters Sep-Pak tC18 Short cartridge, followed by formulation in a mixture of sterile saline and ethanol (10:1 v/v), with total volume of ca. 13 ml. The resulting solution was then filtered through a 0.22 µm sterile filter.

*Metabolite analysis* Plasma was isolated by centrifugation at 4000 rpm for 2 minutes. The resulting plasma was then combined with acetonitrile to precipitate proteins and centrifuged at 6000 rpm for 4 minutes. The supernatant was subsequently diluted with 2 mL of water and analyzed using a radio-HPLC system to determine the unchanged tracer fraction. The amount of radioactivity in blood, in plasma and left in precipitate was measured separately with a NaI(Tl) well counter (PerkinElmer 2480 Automatic Gamma Counter Wizard 2). The HPLC system consisted of an Agilent binary pump (Agilent 1200 series) connected to a manual injection valve (Rheodyne 7725i), a 5.0 mL loop, and a radiation detector (Oyokoken S-2493Z) housed within a 50 mm lead shield. Data collection and control of the HPLC system were performed using chromatographic software (EZChrom Edition, Version A.04.05, Agilnet). Chromatographic separation was performed on a semi-preparative reversed-phase ACE C18 HL column (5 µm, 250 × 10 mm) using gradient elution. The mobile phase consisted of acetonitrile (solvent A) and 0.1 M ammonium formate (solvent B), delivered at a flow rate of 5.0 mL/min. The following program was followed: 0–4.0 min, (A/B) 40:60-90:10 v/v; 4.0–6.0 min, (A/B) 90:10 v/v; 6–6.1 min, (A/B) 90:10-40:60 v/v. The area of each obtained peak in the radiochromatogram was expressed as a percentage of the sum of the areas of all detected radioactive peaks (with decay correction).

*Solena* For every subject, 1 T1-weighted image was used. The image was realigned, cropped and segmented. Freesurfer version 6.0.0 was run on the cropped MRI, with an additional "-brainstem-structures" Freesurfer segmentation[1]. Composite ROIs were created based on the cortical subregions found in table S2.

For each of the 2 PET runs found per subject, the first 4 minutes of the measurement was used as a reference for realignment and motion correction, using the realignment function from Statistical Parametric Mapping (SPM12). The motion-corrected PET was co-registered to the cropped MRI and TACs were extracted. Volumes were extracted from the ROIs in PET space.

*PICNIC* The BIDS app PICNIC (https://github.com/bil-mind-nyspi/PICNIC) was run as a docker container (version 0.1.1, downloadable with docker pull mfschmidt/picnic:0.1.1) with an input deck for each individual. The input deck specifies the file paths to the MRI and PET to be imported as well as the steps to be run.

For each subject, one T1-weighted image was used and FreeSurfer 7.3.2 was run. All outputs were reoriented, and bilateral ROIs were generated, as well as whole brain, gray matter, white matter, subcortical and ventricle masks.

For each of the 2 PET runs found per subject, the following preprocessing steps were performed. First, motion correction was performed using the following parameters: Type=mcflirt, Ref Vol=8, Smooth=4, Crop Start=4. The McFLIRT coregistration is based on the McFLIRT coregestration in FSL[2]. Next, coregistration of the PET images to the MRI with Coregistration Automated Multi Run Approach (CAMRA) was performed using the freesurfer output (T1, whole brain mask, grey matter and white matter masks) and the following parameters: Type=LCF, DOF=6, Cost=mutualinfo. This coregistration using the Lowest Cost Function (LCF) is based on FLIRT in FSL. Lastly, using the freesurfer bilateral_wmparc the TACs were extracted.

*PETPrep* The boilerplate generated by running PETPrep, as run with the "- -ignore flair" tag and with including FLAIR scans is included in the next three paragraphs. This boilerplate is generated by PETPrep and is copied here verbatim from the HTML-reports.

Results included in this manuscript come from preprocessing performed using *PETPrep* 0.0.2 (Esteban et al.[3]; Esteban et al.[4]; RRID:SCR_016216), which is based on *Nipype* 1.9.2 (Gorgolewski et al.[5]; Gorgolewski et al.[6]; RRID:SCR_002502).

*Anatomical data preprocessing* A total of 1 T1-weighted (T1w) images were found within the input BIDS dataset. The T1w image was corrected for intensity non-uniformity (INU) with `N4BiasFieldCorrection`[7], distributed with ANTs 2.6.2[8] RRID:SCR_004757, and used as T1w-reference throughout the workflow. The T1w-reference was then skull-stripped with a *Nipype* implementation of the `antsBrainExtraction.sh` workflow (from ANTs), using OASIS30ANTs as target template. Brain tissue segmentation of cerebrospinal fluid (CSF), white-matter (WM) and gray-matter (GM) was performed on the brain-extracted T1w using `fast` FSL (version unknown), RRID:SCR_002823,[9]. Brain surfaces were reconstructed using `recon-all` FreeSurfer 7.4.1, RRID:SCR_001847,[10], and the brain mask estimated previously was refined with a custom variation of the method to reconcile ANTs-derived and FreeSurfer-derived segmentations of the cortical gray-matter of Mindboggle RRID:SCR_002438,[11]. Volume-based spatial normalization to one standard space (MNI152NLin2009cAsym) was performed through nonlinear registration with `antsRegistration` (ANTs 2.6.2), using brain-extracted versions of both T1w reference and the T1w template. The following template was were selected for spatial normalization and accessed with *TemplateFlow* 24.2.2,[12]: *ICBM 152 Nonlinear Asymmetrical template version 2009c* [Fonov et al.[13], RRID:SCR_008796; TemplateFlow ID: MNI152NLin2009cAsym].

*Anatomical data preprocessing with FLAIR* A total of 1 T1-weighted (T1w) images were found within the input BIDS dataset. The T1w image was corrected for intensity non-uniformity (INU) with `N4BiasFieldCorrection`[7], distributed with ANTs 2.6.2[8] RRID:SCR_004757, and used as T1w-reference throughout the workflow. The T1w-reference was then skull-stripped with a *Nipype* implementation of the `antsBrainExtraction.sh` workflow (from ANTs), using OASIS30ANTs as target template. Brain tissue segmentation of cerebrospinal fluid (CSF), white-matter (WM) and gray-matter (GM) was performed on the brain-extracted T1w using `fast` FSL (version unknown), RRID:SCR_002823,[9]. Brain surfaces were reconstructed using `recon-all` FreeSurfer 7.4.1, RRID:SCR_001847,[10], and the brain mask estimated previously was refined with a custom variation of the method to reconcile ANTs-derived and FreeSurfer-derived segmentations of the cortical gray-matter of Mindboggle RRID:SCR_002438,[11]. A FLAIR image was used to improve pial surface refinement. Brain surfaces were reconstructed using `recon-all` FreeSurfer 7.4.1, RRID:SCR_001847,[10], and the brain mask estimated previously was refined with a custom variation of the method to reconcile ANTs-derived and FreeSurfer-derived segmentations of the cortical gray-matter of Mindboggle RRID:SCR_002438,[11]. Volume-based spatial normalization to one standard space (MNI152NLin2009cAsym) was performed through nonlinear registration with `antsRegistration` (ANTs 2.6.2), using brain-extracted versions of both T1w reference and the T1w template. The following template was were selected for spatial normalization and accessed with *TemplateFlow* 24.2.2,[12]: *ICBM 152 Nonlinear Asymmetrical template version 2009c* [Fonov et al.[13], RRID:SCR_008796; TemplateFlow ID: MNI152NLin2009cAsym].

*PET data preprocessing* For each of the 2 PET runs found per subject (across all tasks and sessions), the following preprocessing steps were performed. Robust head motion estimation and correction were carried out after generating a reference image, which was subsequently coregistered to the T1-weighted anatomical image. A brain mask was computed and the structural image was segmented with the `gtm` segmentation workflow from FreeSurfer. Head-motion parameters with respect to the PET reference (transformation matrices, and six corresponding rotation and translation parameters) are estimated before any spatiotemporal filtering using FreeSurfer's `mri_robust_template`. Several confounding time-series were calculated based on the *preprocessed PET*: framewise displacement (FD), DVARS and three region-wise global signals. FD was computed using two formulations following Power (absolute sum of relative motions, Power et al.[14]) and Jenkinson (relative root mean square displacement between affines, Jenkinson et al.[15]). FD and DVARS are calculated for each PET run, both using their implementations in *Nipype* following the definitions by[14]. The three global signals are extracted within the CSF, the WM, and the whole-brain masks. Additionally, a set of physiological regressors were extracted to allow for component-based noise correction *CompCor*,[16]. Principal components are estimated after high-pass filtering the *preprocessed PET* time-series (using a discrete cosine filter with 128s cut-off) for the two *CompCor* variants: temporal (tCompCor) and anatomical (aCompCor). tCompCor components are then calculated from the top 2% variable voxels within the brain mask. For aCompCor, three probabilistic masks (CSF, WM and combined CSF+WM) are generated in anatomical space. The implementation differs from that of Behzadi et al. in that instead of eroding the masks by 2 pixels on PET space, a mask of pixels that likely contain a volume fraction of GM is subtracted from the aCompCor masks. This mask is obtained by dilating a GM mask extracted from the FreeSurfer's *aseg* segmentation, and it ensures components are not extracted from voxels containing a minimal fraction of GM. Finally, these masks are resampled into PET space and binarized by thresholding at 0.99 (as in the original implementation). Components are also calculated separately within the WM and CSF masks. For each CompCor decomposition, the $k$ components with the largest singular values are retained, such that the retained components' time series are sufficient to explain 50 percent of variance across the nuisance mask (CSF, WM, combined, or temporal). The remaining components are dropped from consideration. The head-motion estimates calculated in the correction step were also placed within the corresponding confounds file. The confound time series derived from head motion estimates and global signals were expanded with the inclusion of temporal derivatives and quadratic terms for each[17]. Frames that exceeded a threshold of 0.5 mm FD or 1.5 standardized DVARS were annotated as motion outliers. Additional nuisance timeseries are calculated by means of principal components analysis of the signal found within a thin band (*crown*) of voxels around the edge of the brain, as proposed by[18]. All resamplings can be performed with *a single interpolation step* by composing all the pertinent transformations (i.e. head-motion transform matrices, susceptibility distortion correction when available, and co-registrations to anatomical and output spaces). Gridded (volumetric) resamplings were performed using `nitransforms`, configured with cubic B-spline interpolation.

Many internal operations of *PETPrep* use *Nilearn* 0.11.1[19] RRID:SCR_001362, mostly within the PET processing workflow. For more details of the pipeline, see the section corresponding to workflows in *PETPrep*'s documentation.

*Copyright Waiver* The above boilerplate text was automatically generated by PETPrep with the express intention that users should copy and paste this text into their manuscripts *unchanged*. It is released under the CC0 license.

*Bloodstream blood-to-plasma ratio and parent fraction modeling* The blood-to-plasma ratio (BPR) and parent fraction was fit with Bloodstream using the HGAM method, using the following configuration (using both a PET and subject smooth).

```
"ParentFraction": {
  "Method": ["Fit Hierarchically: HGAM"],
  "set_ppf0": [true],
  "starttime": [0],
  "endtime": ["Inf"],
  "gam_k": ["6"],
  "hgam_formula": ["s(log(time)) + s(log(time), pet, bs='fs', k=6) +
  s(log(time), sub, bs='fs', k =6)"]
},
"BPR": {
  "Method": ["Fit Hierarchically: HGAM"],
  "starttime": [0],
  "endtime": ["Inf"],
  "gam_k": [6],
  "hgam_formula": ["s(time, k=6) + s(time, pet, bs='fs', k=6) +
  s(time, sub, bs='fs', k =6)"]
}
```

**Table S1.** Acquisition parameters of the different structural MRI sequences. Time to echo (TE) and Time to Repetition (TR) both in ms. No FLAIR MRI was collected using the SIGNA Premier scanner.

| | DISCOVERY MR750 | | SIGNA Premier |
|---|---|---|---|
| Sequence | FLAIR | T1 | T1 |
| TE, mean (sd) | 112.68 (1.46) | 2.94 (0) | 2.97 (0) |
| TR, mean (sd) | 7000 (0) | 7.64 (0) | 7.26 (0) |
| Voxel Size (mm) | 0.600x0.527x0.527 | 1.000x1.000x1.000 | 1.000x1.000x1.000 |

**Table S2.** All subregions used to define the regions of interest for the three different PET preprocessing tools. PICNIC and Solena have bilaterally defined regions, whereas PETPrep specifies most regions for the left hemisphere (lh) and right hemisphere (rh) separately.

| PICNIC | PETPrep | | Solena |
|---|---|---|---|
| bilateral | lh | rh | |
| Frontal cortex | | | |
| ctx-bilateral-parstriangularis | ctx-lh-parstriangularis | ctx-rh-parstriangularis | ctx-parstriangularis |
| ctx-bilateral-rostralmiddlefrontal | ctx-lh-rostralmiddlefrontal | ctx-rh-rostralmiddlefrontal | ctx-rostralmiddlefrontal |
| ctx-bilateral-caudalmiddlefrontal | ctx-lh-caudalmiddlefrontal | ctx-rh-caudalmiddlefrontal | ctx-caudalmiddlefrontal |
| ctx-bilateral-frontalpole | ctx-lh-frontalpole | ctx-rh-frontalpole | ctx-frontalpole |
| ctx-bilateral-medialorbitofrontal | ctx-lh-medialorbitofrontal | ctx-rh-medialorbitofrontal | ctx-medialorbitofrontal |
| ctx-bilateral-superiorfrontal | ctx-lh-superiorfrontal | ctx-rh-superiorfrontal | ctx-superiorfrontal |
| ctx-bilateral-lateralorbitofrontal | ctx-lh-lateralorbitofrontal | ctx-rh-lateralorbitofrontal | ctx-lateralorbitofrontal |
| ctx-bilateral-parsorbitalis | ctx-lh-parsorbitalis | ctx-rh-parsorbitalis | ctx-parsorbitalis |
| Temporal cortex | | | |

**Table S2.** Table S2 (Continued)

| PICNIC | PETPrep | | Solena |
|---|---|---|---|
| ctx-bilateral-inferiortemporal | ctx-lh-inferiortemporal | ctx-rh-inferiortemporal | ctx-inferiortemporal |
| ctx-bilateral-middletemporal | ctx-lh-middletemporal | ctx-rh-middletemporal | ctx-middletemporal |
| ctx-bilateral-superiortemporal | ctx-lh-superiortemporal | ctx-rh-superiortemporal | ctx-superiortemporal |
| ctx-bilateral-transversetemporal | ctx-lh-transversetemporal | ctx-rh-transversetemporal | ctx-transversetemporal |
| ctx-bilateral-entorhinal | ctx-lh-entorhinal | ctx-rh-entorhinal | ctx-entorhinal |
| ctx-bilateral-fusiform | ctx-lh-fusiform | ctx-rh-fusiform | ctx-fusiform |
| ctx-bilateral-temporalpole | ctx-lh-temporalpole | ctx-rh-temporalpole | ctx-temporalpole |
| Occipital cortex | | | |
| ctx-bilateral-lateraloccipital | ctx-lh-lateraloccipital | ctx-rh-lateraloccipital | ctx-lateraloccipital |
| ctx-bilateral-cuneus | ctx-lh-cuneus | ctx-rh-cuneus | ctx-cuneus |
| ctx-bilateral-lingual | ctx-lh-lingual | ctx-rh-lingual | ctx-lingual |
| ctx-bilateral-pericalcarine | ctx-lh-pericalcarine | ctx-rh-pericalcarine | ctx-pericalcarine |
| Parietal cortex | | | |
| ctx-bilateral-inferiorparietal | ctx-lh-inferiorparietal | ctx-rh-inferiorparietal | ctx-inferiorparietal |
| ctx-bilateral-superiorparietal | ctx-lh-superiorparietal | ctx-rh-superiorparietal | ctx-superiorparietal |
| ctx-bilateral-supramarginal | ctx-lh-supramarginal | ctx-rh-supramarginal | ctx-supramarginal |
| ctx-bilateral-precuneus | ctx-lh-precuneus | ctx-rh-precuneus | ctx-precuneus |
| Striatum | | | |
| bilateral-caudate | Left-Caudate | Right-Caudate | Caudate |
| bilateral-accumbens-area | Left-Accumbens-area | Right-Accumbens-area | Accumbens-area |
| bilateral-putamen | Left-Putamen | Right-Putamen | Putamen |
| | | | caudate-intensity-abnormality |
| | | | accumbens-intensity-abnormality |
| | | | putamen-intensity-abnormality |
| Thalamus | | | |
| bilateral-thalamus-proper* | Left-Thalamus | Right-Thalamus | Thalamus |
| | | | Thalamus-Proper |
| | | | thalamus-intensity-abnormality |
| Brainstem | | | |
| brain-stem | Brain-Stem | | Brain-Stem |
| | | | Medulla |
| | | | Midbrain |
| | | | Pons |
| | | | SCP |
| Cerebellar Cortex | | | |
| bilateral-cerebellum-cortex | Left-Cerebellum-Cortex | Right-Cerebellum-Cortex | Cerebellum-Cortex |
| Hippocampus | | | |
| bilateral-hippocampus | Left-Hippocampus | Right-Hippocampus | Hippocampus |
| | | | hippocampus-intensity-abnormality |

### *Supplementary results*

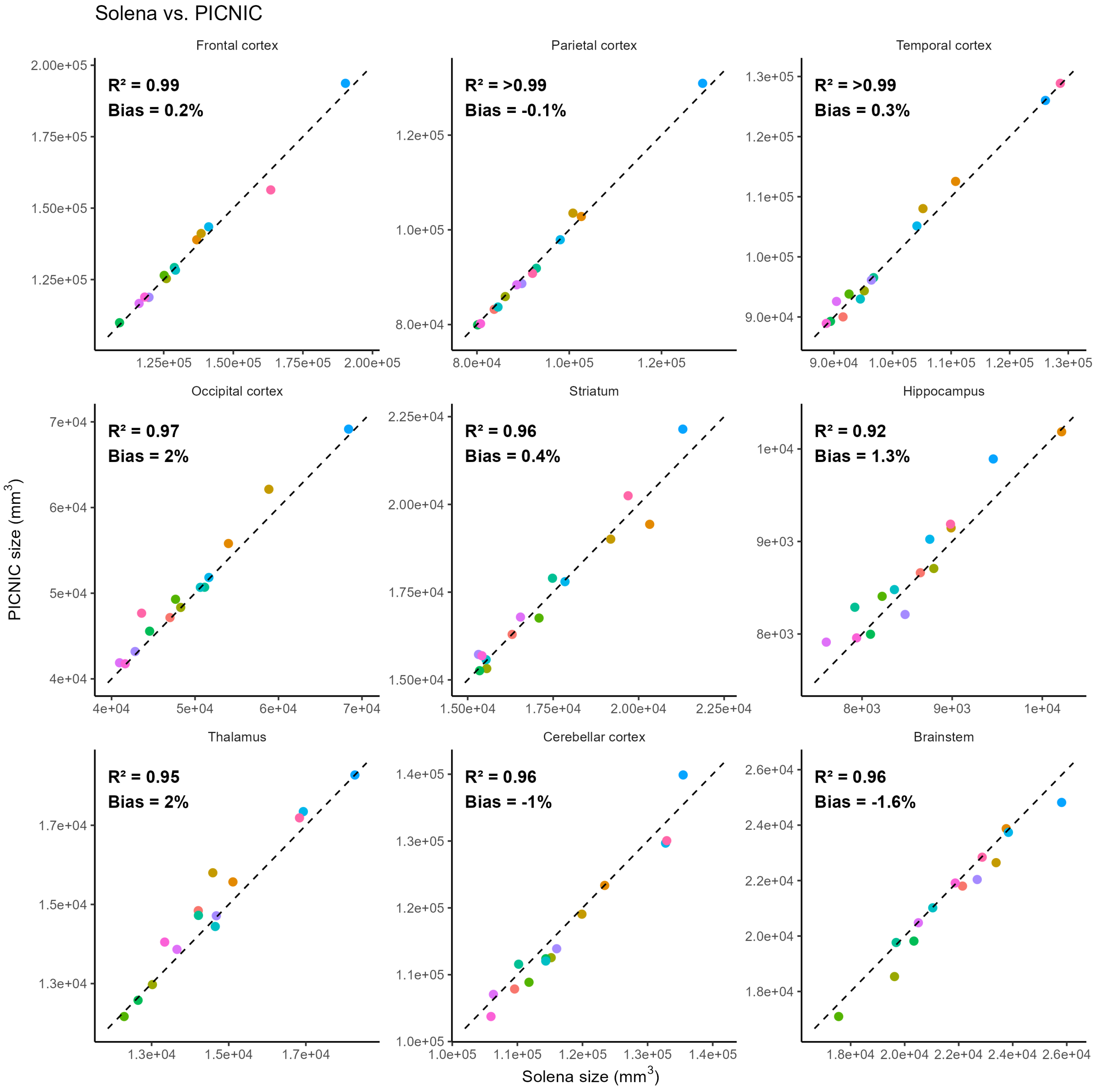


**Figure S1.** Comparison of volumes derived from Solena and PICNIC for the 9 different ROIs. An average volume from the two sessions is used for Solena. Bias is calculated using the PD formula, with positive bias indicating higher volumes for PICNIC.

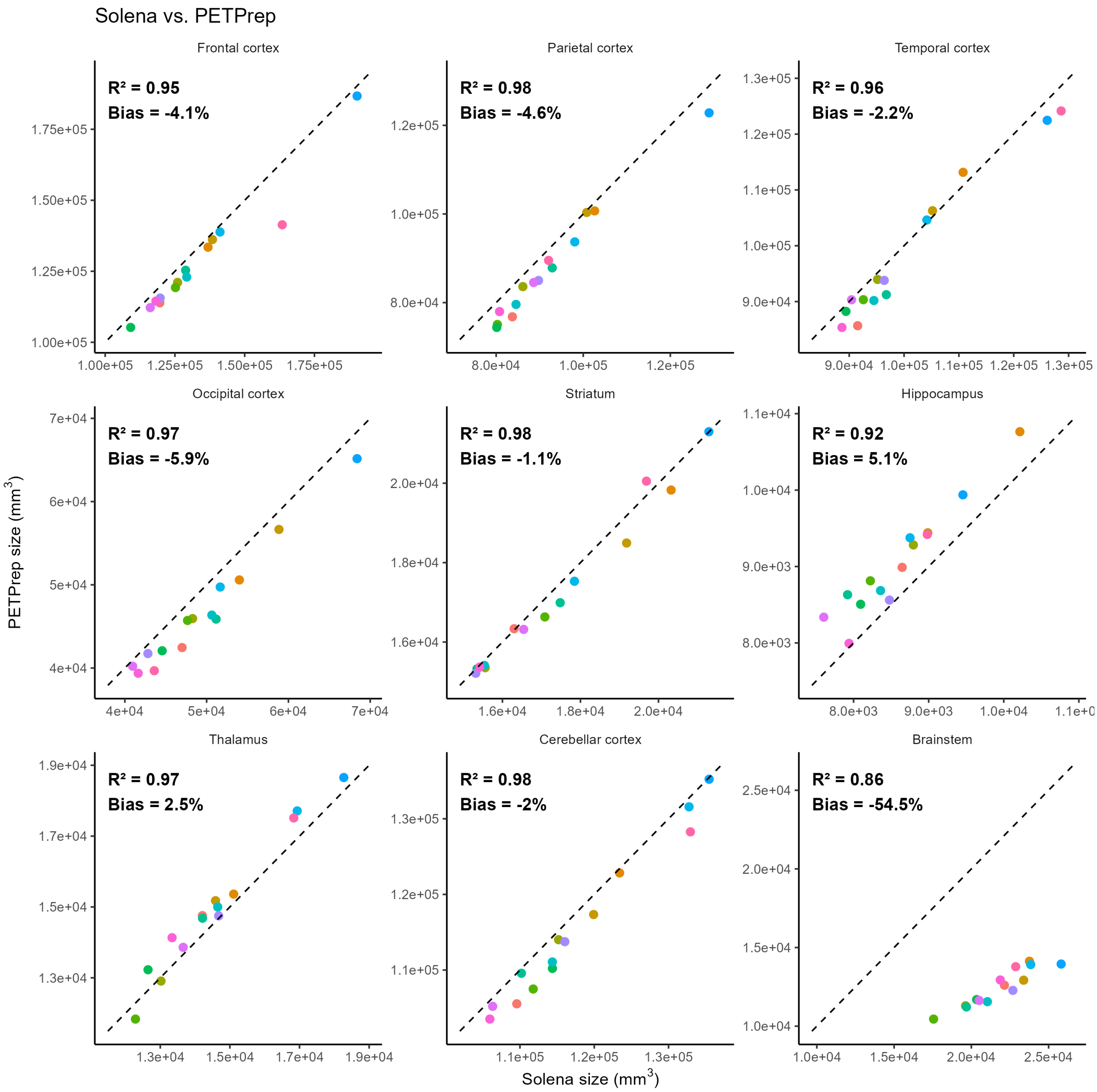


**Figure S2.** Comparison of volumes derived from Solena and PETPrep for the 9 different ROIs. An average volume from the two sessions is used for Solena. Bias is calculated using the PD formula, with positive bias indicating higher volumes for PETPrep.

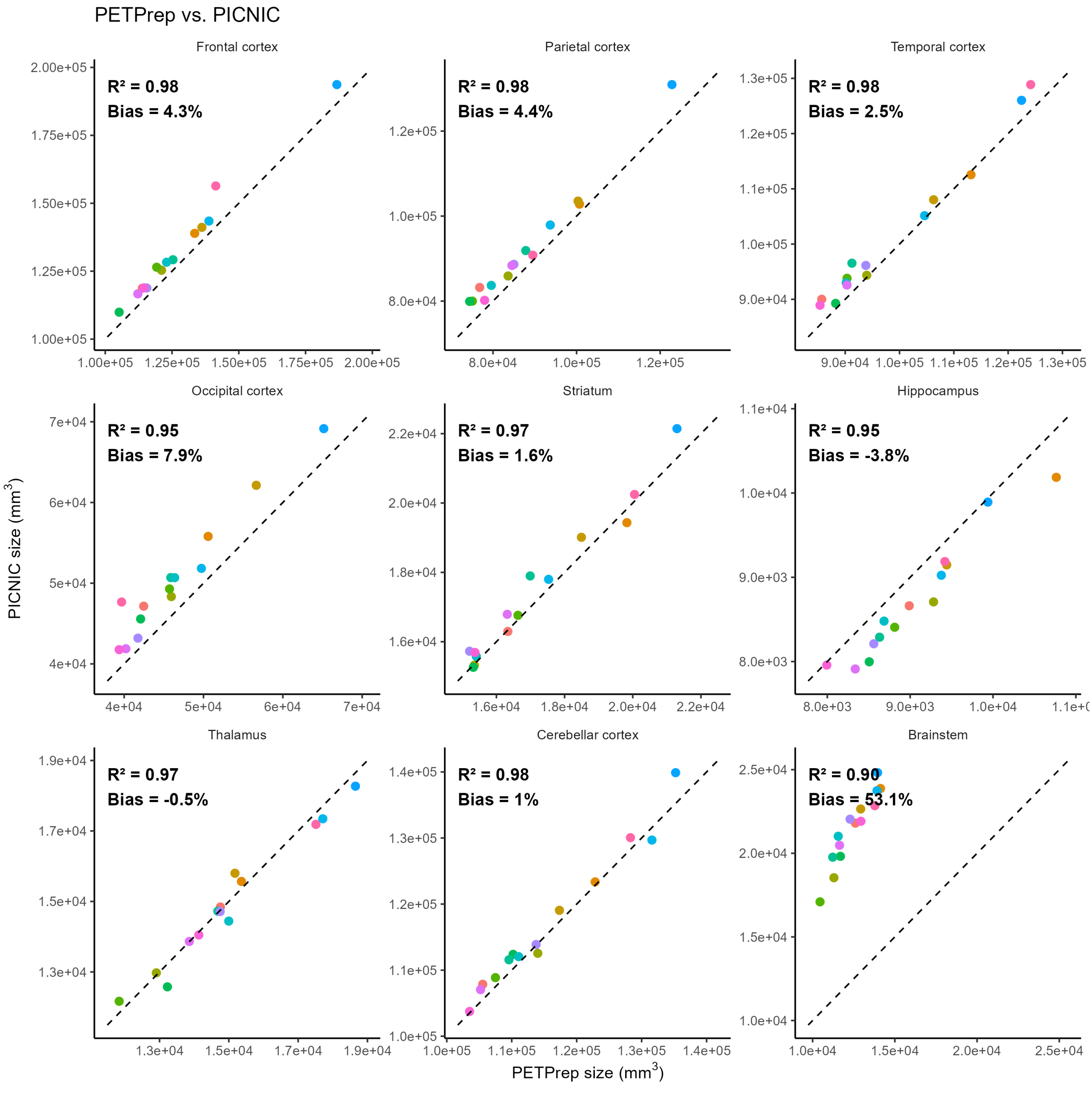


**Figure S3.** Comparison of volumes derived from PETPrep and PICNIC for the 9 different ROIs. Bias is calculated using the PD formula, with positive bias indicating higher volumes for PICNIC.

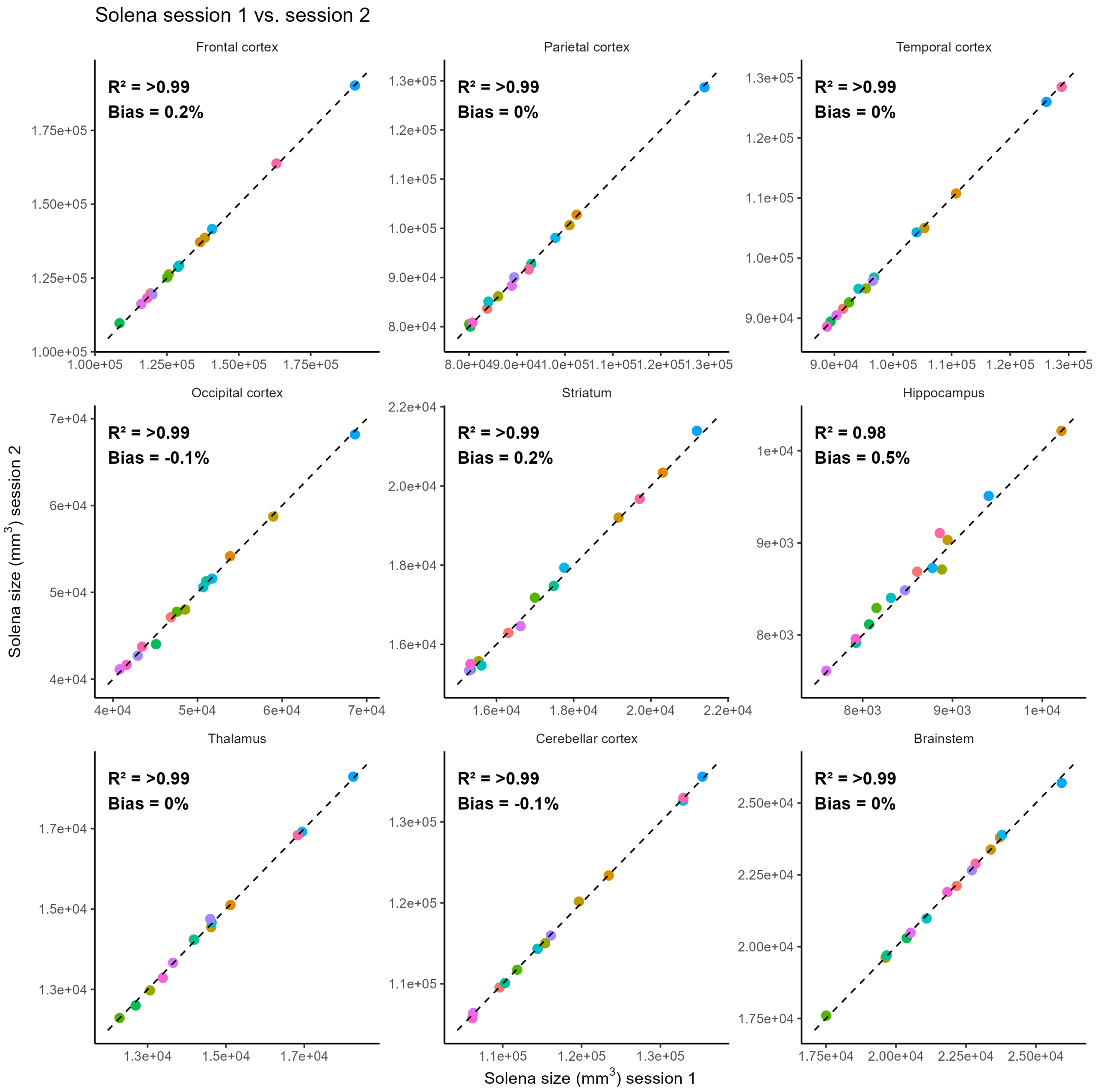


**Figure S4.** Comparison of volumes derived from Solena for the two sessions for the 9 different ROIs. Bias is calculated using the PD formula, with positive bias indicating higher volumes for session 2.

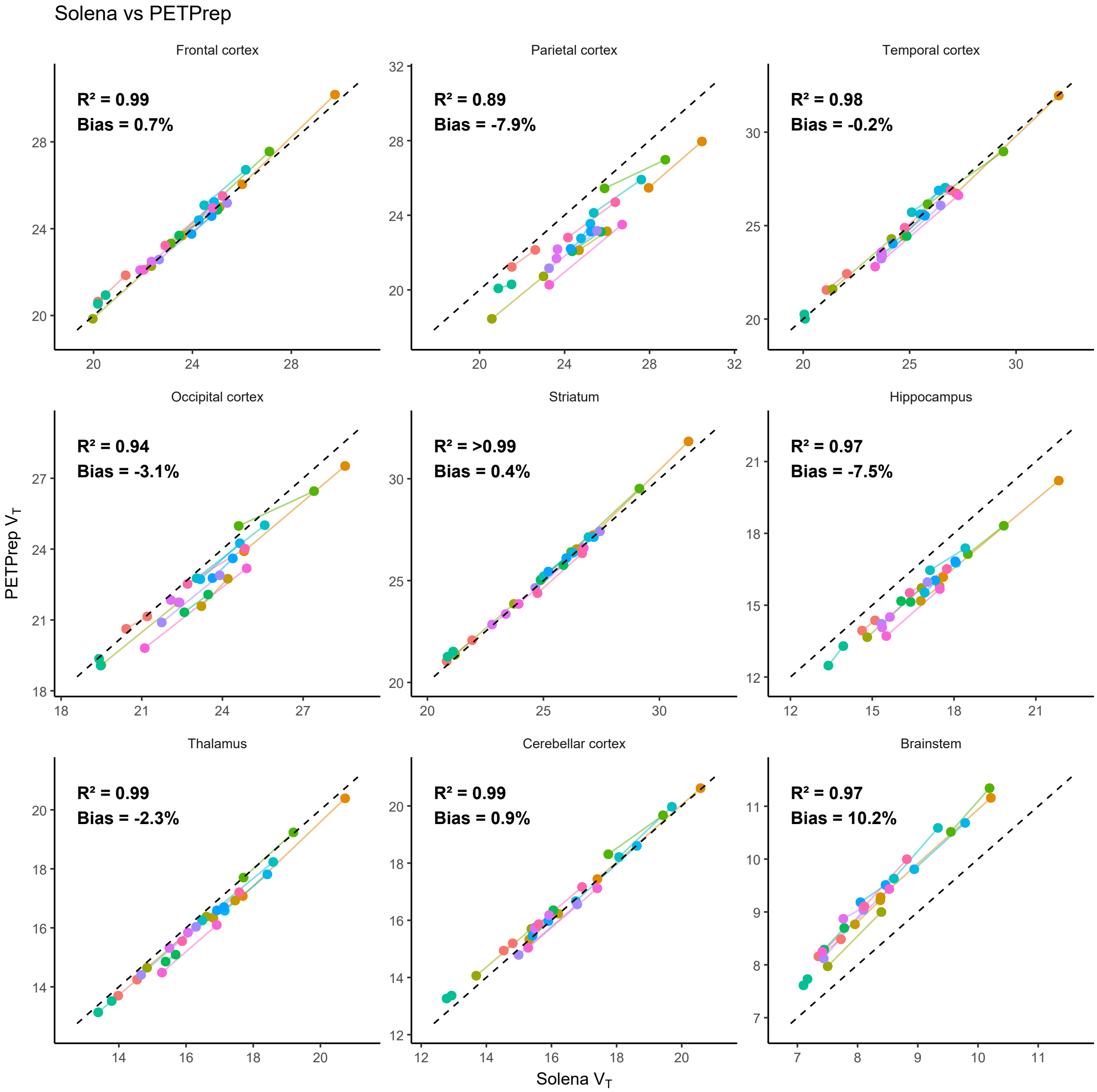


**Figure S5.** Direct comparison of $V_{\mathrm{T}}$ as derived by Solena and PETPrep for the 9 ROIs. Bias is calculated using the PD formula, with positive bias indicating higher $V_{\mathrm{T}}$ for PETPrep.

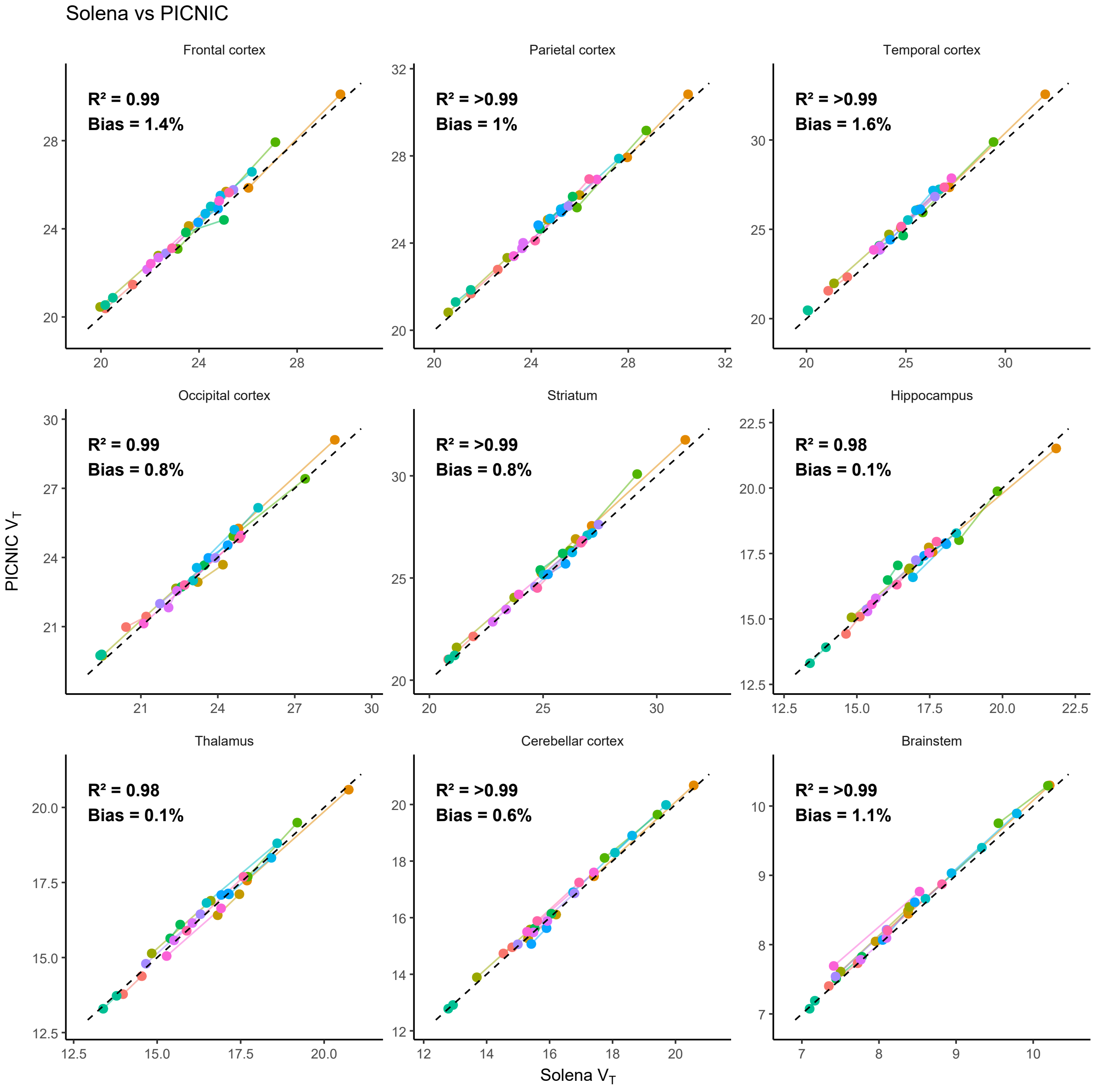


**Figure S6.** Direct comparison of $V_{\mathrm{T}}$ as derived by Solena and PICNIC for the 9 ROIs. Bias is calculated using the PD formula, with positive bias indicating higher $V_{\mathrm{T}}$ for PICNIC.

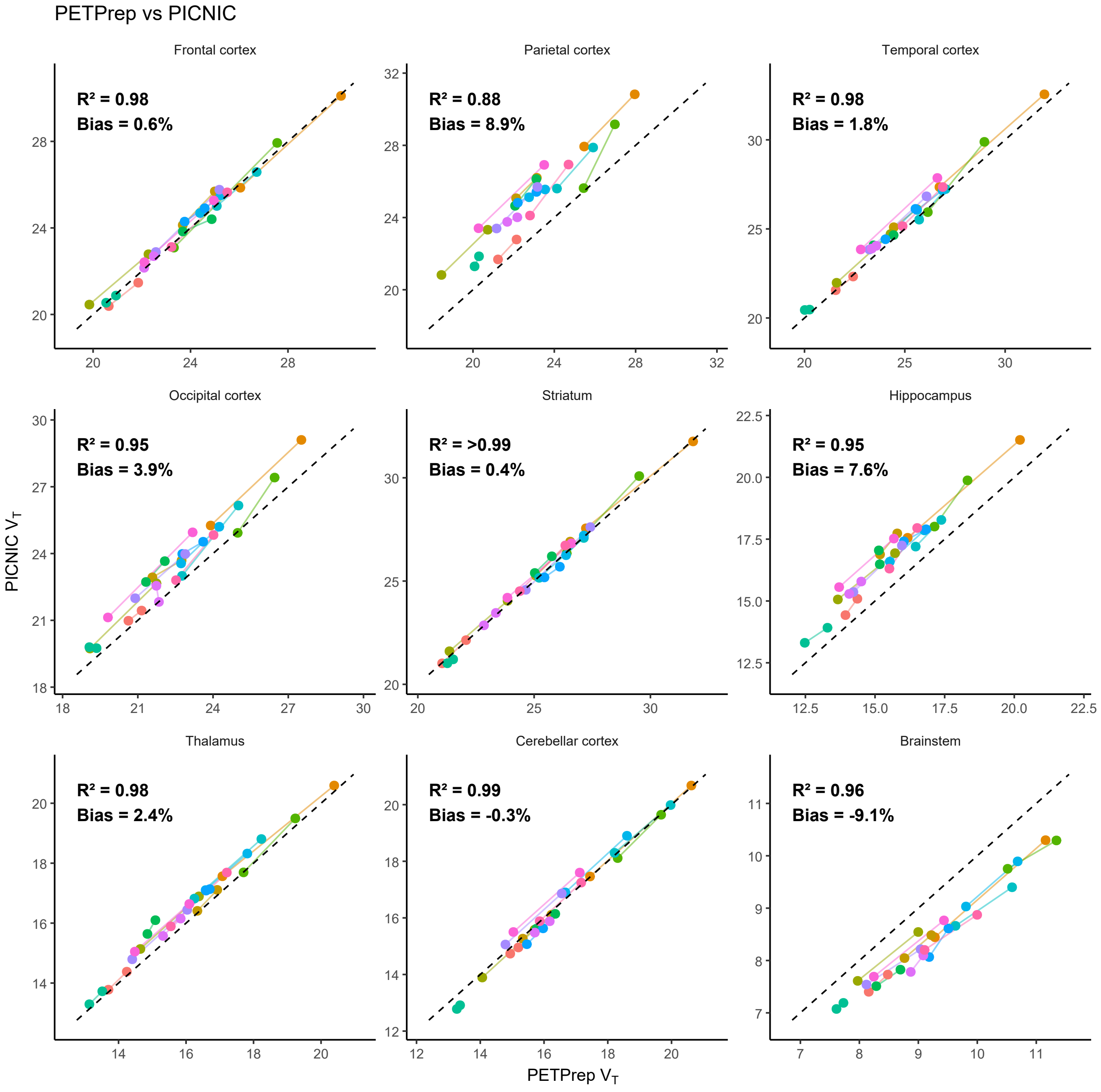


**Figure S7.** Direct comparison of $V_\mathrm{T}$ as derived by PETPrep and PICNIC for the 9 ROIs. Bias is calculated using the PD formula, with positive bias indicating higher $V_\mathrm{T}$ for PICNIC

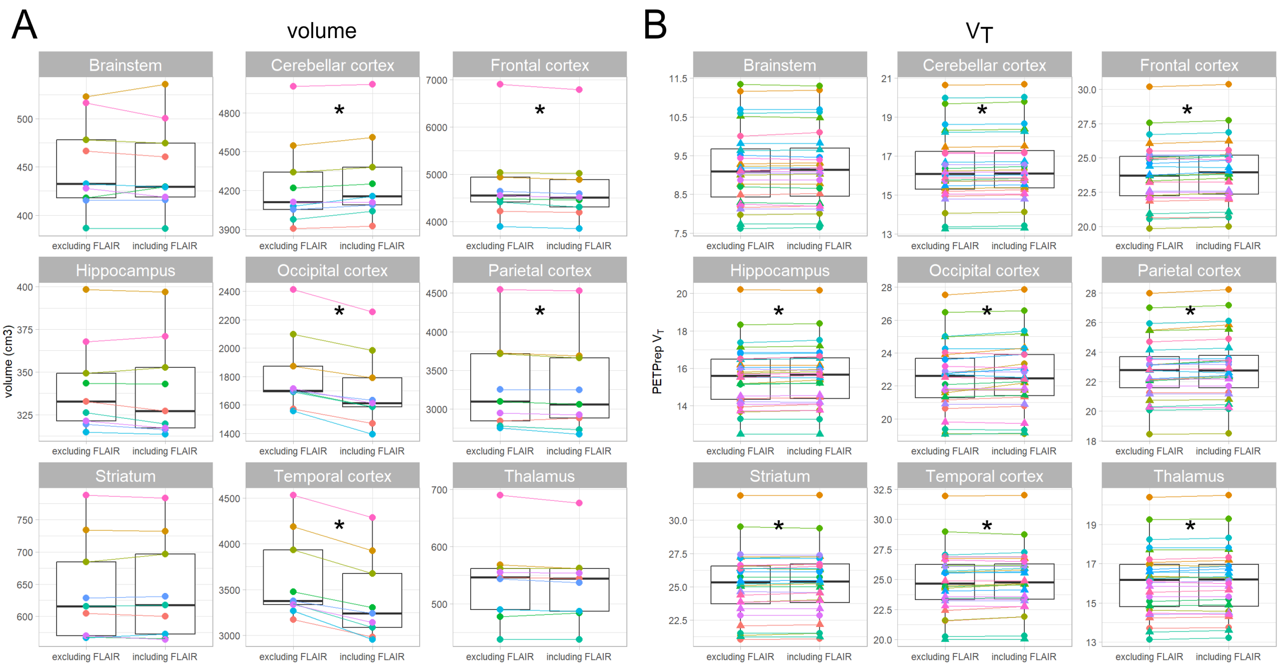


**Figure S8.** Comparison of volumes (A) and $V_{\mathrm{T}}$ values (B) as derived from preprocessing in PETPrep with or without usage of FLAIR. Results only for the subjects with FLAIR scans in the dataset (N=9). For $V_{\mathrm{T}}$ both the first and second session are included (N=18). Comparisons with an uncorrected p-value $< 0.05$ are indicated with *. Data points are color-coded by subject.

**Table S3.** Comparison of $V_{\mathrm{T}}$ as calculated by PETPrep run without or with FLAIR included for 9 subjects, both sessions. $P$-values for paired $t$-tests with N=18. Percentage change calculated as (mean difference / mean excluding FLAIR )*100 %

| Region | Excluding FLAIR, mean (sd) | Including FLAIR, mean (sd) | Difference, mean (95% CI) | Percentage change | p-value |
|---|---|---|---|---|---|
| Frontal cortex | 23.92 (2.70) | 24.09 (2.72) | 0.18 (0.15–0.21) | 0.7% | 6.37e-10* |
| Parietal cortex | 23.03 (2.54) | 23.21 (2.59) | 0.18 (0.13–0.23) | 0.8% | 9.15e-07* |
| Temporal cortex | 24.68 (3.01) | 24.82 (2.95) | 0.14 (0.07–0.21) | 0.6% | 5.94e-04* |
| Occipital cortex | 22.54 (2.40) | 22.77 (2.49) | 0.23 (0.13–0.32) | 1.0% | 7.32e-05* |
| Striatum | 25.13 (2.95) | 25.18 (2.93) | 0.05 (0.00–0.10) | 0.2% | 4.14e-02* |
| Hippocampus | 15.74 (1.88) | 15.80 (1.88) | 0.06 (0.03–0.09) | 0.4% | 1.29e-03* |
| Thalamus | 16.17 (1.99) | 16.22 (2.00) | 0.05 (0.02–0.09) | 0.3% | 7.51e-03* |
| Cerebellar cortex | 16.44 (2.18) | 16.50 (2.18) | 0.06 (0.05–0.08) | 0.4% | 4.35e-09* |
| Brainstem | 9.17 (1.12) | 9.18 (1.12) | 0.01 (-0.01–0.02) | 0.1% | 3.62e-01 |

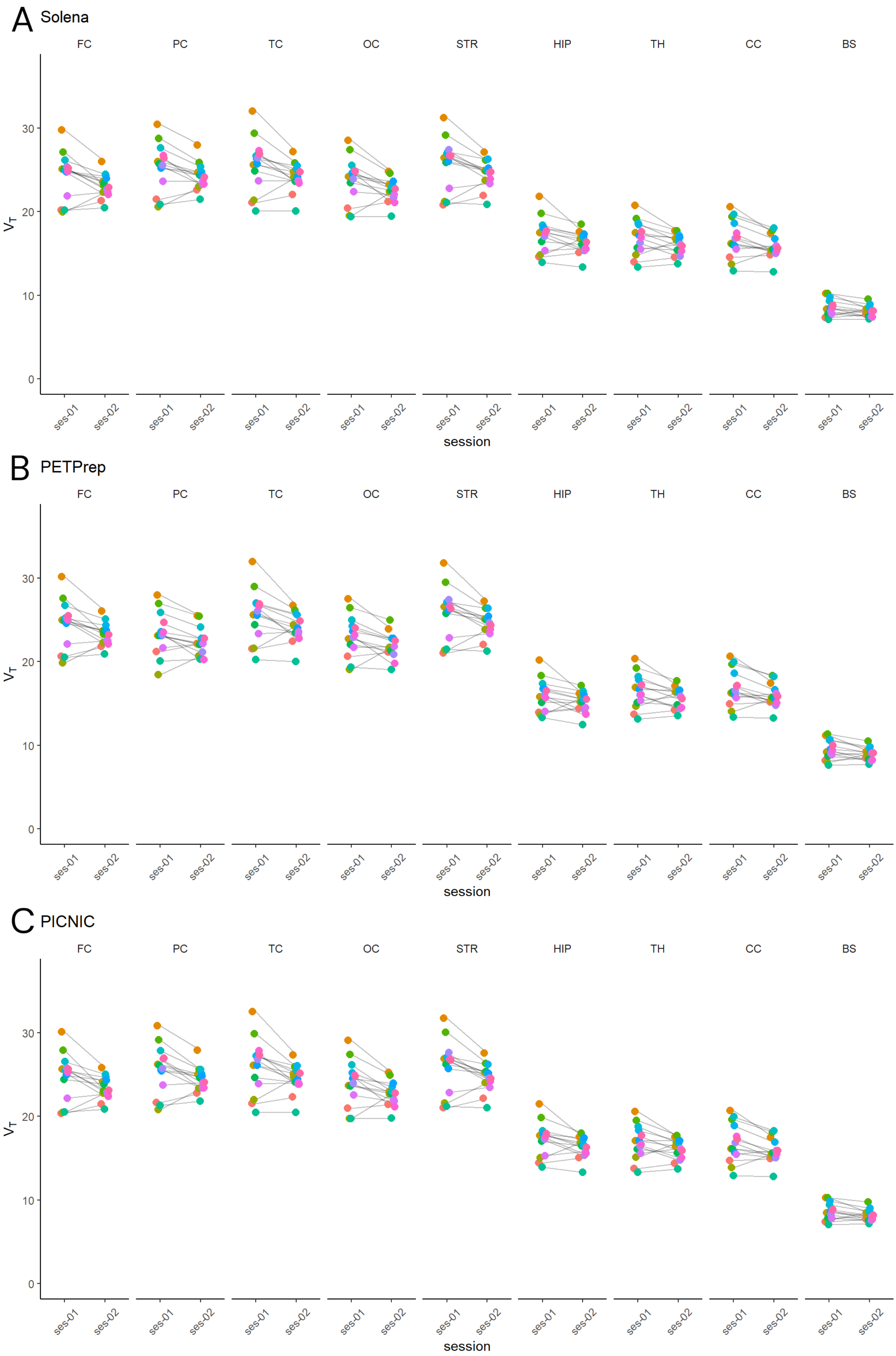


**Figure S9.** $V_{\mathrm{T}}$ for the 9 different ROIs with preprocessing performed by Solena, PETPrep and PICNIC. FC: Frontal cortex, PC: Parietal cortex, TC: Temporal cortex, OC: Occipital cortex, STR: striatum, HIP: Hippocampus, TH: Thalamus, CC: Cerebellar cortex, BS: Brainstem.

**Table S4.** Comparison of $V_T$ as derived using Solena for session 1 and session 2, difference (session 2 - session 1), APD and PD, reported as mean (sd). $ICC_a$ calculated with two-way mixed effects and absolute agreement and reported as estimate (95% CI). P-values are calculated using a paired two-tailed t-test. P-values $< 0.05$ are indicated with *.

| Region | session 1 $V_T$ | session 2 $V_T$ | difference | p-value | APD | PD | $ICC_a$ |
|---|---|---|---|---|---|---|---|
| Frontal cortex | 24.32 (2.83) | 23.07 (1.40) | -1.25 | 0.025* | 7.69 (4.59) | -4.80 (7.74) | 0.58 (0.09–0.84) |
| Parietal cortex | 25.30 (2.87) | 24.20 (1.57) | -1.10 | 0.029* | 6.78 (3.89) | -4.02 (6.87) | 0.67 (0.20–0.88) |
| Temporal cortex | 25.54 (3.23) | 24.15 (1.70) | -1.40 | 0.022* | 7.49 (5.31) | -5.10 (7.79) | 0.62 (0.12–0.86) |
| Occipital cortex | 23.83 (2.68) | 22.55 (1.41) | -1.28 | 0.02* | 7.62 (5.08) | -5.08 (7.79) | 0.56 (0.06–0.83) |
| Striatum | 25.68 (3.11) | 24.49 (1.67) | -1.19 | 0.027* | 6.98 (4.02) | -4.27 (7.00) | 0.68 (0.21–0.89) |
| Hippocampus | 17.22 (2.13) | 16.32 (1.27) | -0.90 | 0.033* | 7.47 (5.31) | -4.95 (7.87) | 0.61 (0.13–0.86) |
| Thalamus | 16.84 (2.05) | 16.07 (1.20) | -0.77 | 0.046* | 7.32 (4.49) | -4.23 (7.65) | 0.65 (0.20–0.87) |
| Cerebellar cortex | 16.73 (2.26) | 15.79 (1.36) | -0.95 | 0.015* | 7.64 (4.75) | -5.31 (7.43) | 0.70 (0.19–0.90) |
| Brainstem | 8.52 (1.03) | 8.09 (0.66) | -0.43 | 0.035* | 7.87 (4.72) | -4.82 (8.00) | 0.62 (0.15–0.86) |

**Table S5.** Comparison of $V_T$ as derived using PETPrep for session 1 and session 2, difference (session 2 - session 1), APD and PD, reported as mean (sd). $ICC_a$ calculated with two-way mixed effects and absolute agreement and reported as estimate (95% CI). P-values are calculated using a paired two-tailed t-test. P-values $< 0.05$ are indicated with *.

| Region | session 1 $V_T$ | session 2 $V_T$ | difference | p-value | APD | PD | $ICC_a$ |
|---|---|---|---|---|---|---|---|
| Frontal cortex | 24.49 (2.87) | 23.24 (1.35) | -1.25 | 0.031* | 7.72 (4.81) | -4.73 (7.95) | 0.56 (0.07–0.83) |
| Parietal cortex | 23.33 (2.56) | 22.41 (1.65) | -0.91 | 0.036* | 6.42 (3.80) | -3.68 (6.65) | 0.72 (0.28–0.90) |
| Temporal cortex | 25.48 (3.11) | 24.10 (1.74) | -1.39 | 0.021* | 7.52 (5.13) | -5.14 (7.66) | 0.61 (0.11–0.86) |
| Occipital cortex | 23.04 (2.42) | 21.94 (1.53) | -1.11 | 0.022* | 6.91 (4.76) | -4.62 (7.14) | 0.61 (0.12–0.86) |
| Striatum | 25.80 (3.13) | 24.59 (1.64) | -1.21 | 0.029* | 6.90 (4.42) | -4.32 (7.12) | 0.66 (0.19–0.88) |
| Hippocampus | 15.97 (1.92) | 15.16 (1.21) | -0.81 | 0.047* | 7.75 (5.73) | -4.86 (8.49) | 0.57 (0.10–0.84) |
| Thalamus | 16.47 (2.05) | 15.70 (1.22) | -0.77 | 0.049* | 7.37 (4.88) | -4.32 (7.88) | 0.64 (0.19–0.87) |
| Cerebellar cortex | 16.88 (2.18) | 15.95 (1.36) | -0.94 | 0.014* | 7.51 (4.74) | -5.27 (7.30) | 0.69 (0.17–0.89) |
| Brainstem | 9.45 (1.18) | 8.95 (0.75) | -0.50 | 0.024* | 7.91 (4.77) | -5.07 (7.90) | 0.65 (0.16–0.87) |

**Table S6.** Comparison of $V_T$ as derived using PICNIC for session 1 and session 2, difference (session 2 - session 1), APD and PD, reported as mean (sd). $ICC_a$ calculated with two-way mixed effects and absolute agreement and reported as estimate (95% CI). P-values are calculated using a paired two-tailed t-test. P-values $< 0.05$ are indicated with *.

| Region | session 1 $V_T$ | session 2 $V_T$ | difference | p-value | APD | PD | $ICC_a$ |
|---|---|---|---|---|---|---|---|
| Frontal cortex | 24.67 (2.88) | 23.37 (1.36) | -1.30 | 0.033* | 7.77 (5.45) | -4.92 (8.28) | 0.52 (0.04–0.81) |
| Parietal cortex | 25.59 (2.90) | 24.41 (1.50) | -1.18 | 0.032* | 7.14 (4.39) | -4.29 (7.37) | 0.62 (0.14–0.86) |
| Temporal cortex | 25.98 (3.28) | 24.49 (1.68) | -1.49 | 0.021* | 7.63 (5.63) | -5.37 (7.96) | 0.59 (0.09–0.85) |
| Occipital cortex | 24.04 (2.68) | 22.72 (1.46) | -1.32 | 0.017* | 7.57 (5.25) | -5.26 (7.72) | 0.56 (0.05–0.83) |
| Striatum | 25.91 (3.21) | 24.65 (1.68) | -1.27 | 0.026* | 7.14 (4.40) | -4.49 (7.24) | 0.66 (0.18–0.88) |
| Hippocampus | 17.26 (2.07) | 16.32 (1.22) | -0.94 | 0.026* | 7.96 (4.87) | -5.19 (7.93) | 0.58 (0.09–0.85) |
| Thalamus | 16.86 (2.06) | 16.09 (1.23) | -0.78 | 0.045* | 7.45 (4.43) | -4.30 (7.71) | 0.65 (0.19–0.87) |
| Cerebellar cortex | 16.84 (2.33) | 15.88 (1.44) | -0.96 | 0.013* | 7.57 (4.81) | -5.36 (7.34) | 0.71 (0.20–0.90) |
| Brainstem | 8.61 (1.05) | 8.18 (0.68) | -0.43 | 0.036* | 7.81 (4.64) | -4.74 (7.94) | 0.63 (0.17–0.87) |

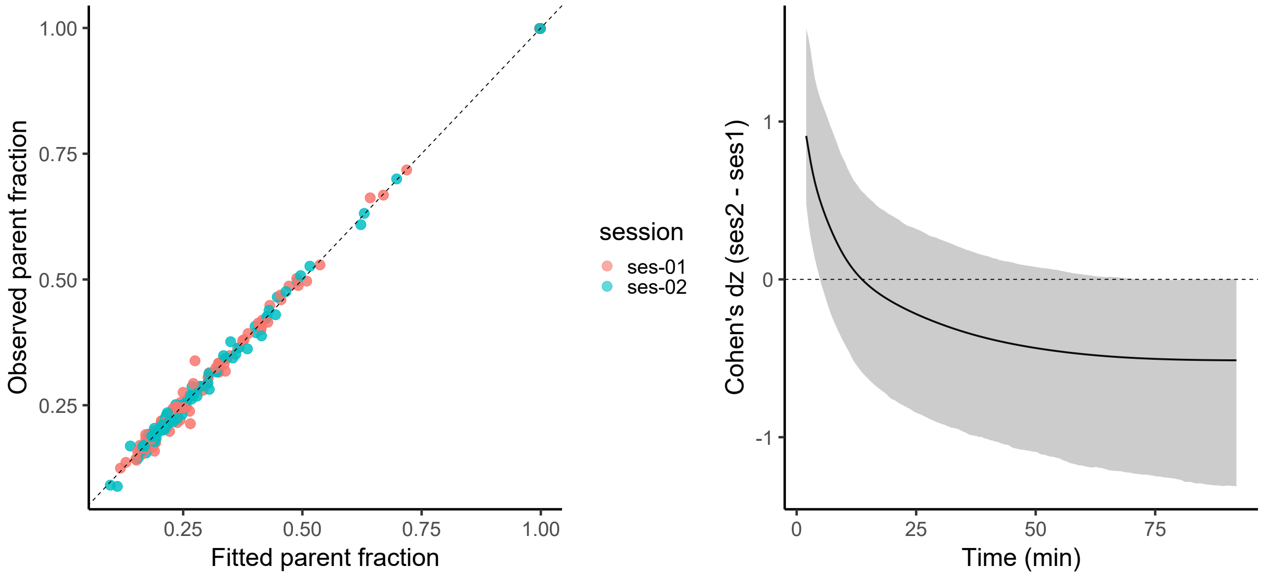


**Figure S10.** Comparison of fitted and observed values for the parent fraction (left) and Cohen's dz (right). Estimated Cohens dz for the fitted parent fraction with the 95% confidence interval. A positive Cohen's dz means fitted values were higher for session 2.

**Table S7.** Proportional standard error ($\times 100\%$) of $V_{\mathrm{T}}$ estimates and difference (ses-02 - ses-01) for both sessions for each tool. P-values are calculated using a paired two-tailed t-test.

| Region | ses-01 | ses-02 | mean difference | p-val |
|---|---|---|---|---|
| Solena | | | | |
| Frontal cortex | 1.132 (0.226) | 1.122 (0.281) | -0.009 | 0.869 |
| Parietal cortex | 1.169 (0.248) | 1.140 (0.278) | -0.029 | 0.682 |
| Temporal cortex | 1.195 (0.235) | 1.129 (0.277) | -0.066 | 0.418 |
| Occipital cortex | 1.096 (0.337) | 1.094 (0.385) | -0.001 | 0.979 |
| Striatum | 1.142 (0.216) | 1.107 (0.268) | -0.035 | 0.635 |
| Hippocampus | 2.542 (0.595) | 2.549 (0.531) | 0.007 | 0.957 |
| Thalamus | 1.568 (0.311) | 1.653 (0.354) | 0.085 | 0.298 |
| Cerebellar cortex | 1.478 (0.339) | 1.557 (0.304) | 0.078 | 0.43 |
| Brainstem | 1.353 (0.312) | 1.391 (0.292) | 0.038 | 0.652 |
| PETPrep | | | | |
| Frontal cortex | 1.044 (0.227) | 1.005 (0.283) | -0.039 | 0.495 |
| Parietal cortex | 1.081 (0.307) | 1.048 (0.272) | -0.033 | 0.699 |
| Temporal cortex | 1.151 (0.240) | 1.078 (0.266) | -0.073 | 0.368 |
| Occipital cortex | 1.021 (0.328) | 1.006 (0.321) | -0.015 | 0.835 |
| Striatum | 1.080 (0.202) | 1.057 (0.245) | -0.022 | 0.754 |
| Hippocampus | 2.792 (0.622) | 2.789 (0.635) | -0.002 | 0.986 |
| Thalamus | 1.571 (0.313) | 1.666 (0.348) | 0.095 | 0.258 |
| Cerebellar cortex | 1.513 (0.332) | 1.579 (0.322) | 0.066 | 0.508 |
| Brainstem | 1.467 (0.261) | 1.525 (0.309) | 0.058 | 0.523 |
| PICNIC | | | | |
| Frontal cortex | 1.016 (0.124) | 0.990 (0.317) | -0.026 | 0.728 |
| Parietal cortex | 1.018 (0.215) | 1.022 (0.238) | 0.003 | 0.967 |
| Temporal cortex | 1.061 (0.185) | 0.976 (0.260) | -0.085 | 0.308 |
| Occipital cortex | 1.044 (0.202) | 0.989 (0.304) | -0.055 | 0.384 |
| Striatum | 1.081 (0.158) | 1.087 (0.303) | 0.006 | 0.943 |
| Hippocampus | 2.398 (0.574) | 2.436 (0.551) | 0.038 | 0.746 |
| Thalamus | 1.449 (0.285) | 1.575 (0.345) | 0.126 | 0.176 |
| Cerebellar cortex | 1.406 (0.335) | 1.463 (0.313) | 0.057 | 0.582 |
| Brainstem | 1.335 (0.304) | 1.368 (0.306) | 0.033 | 0.725 |

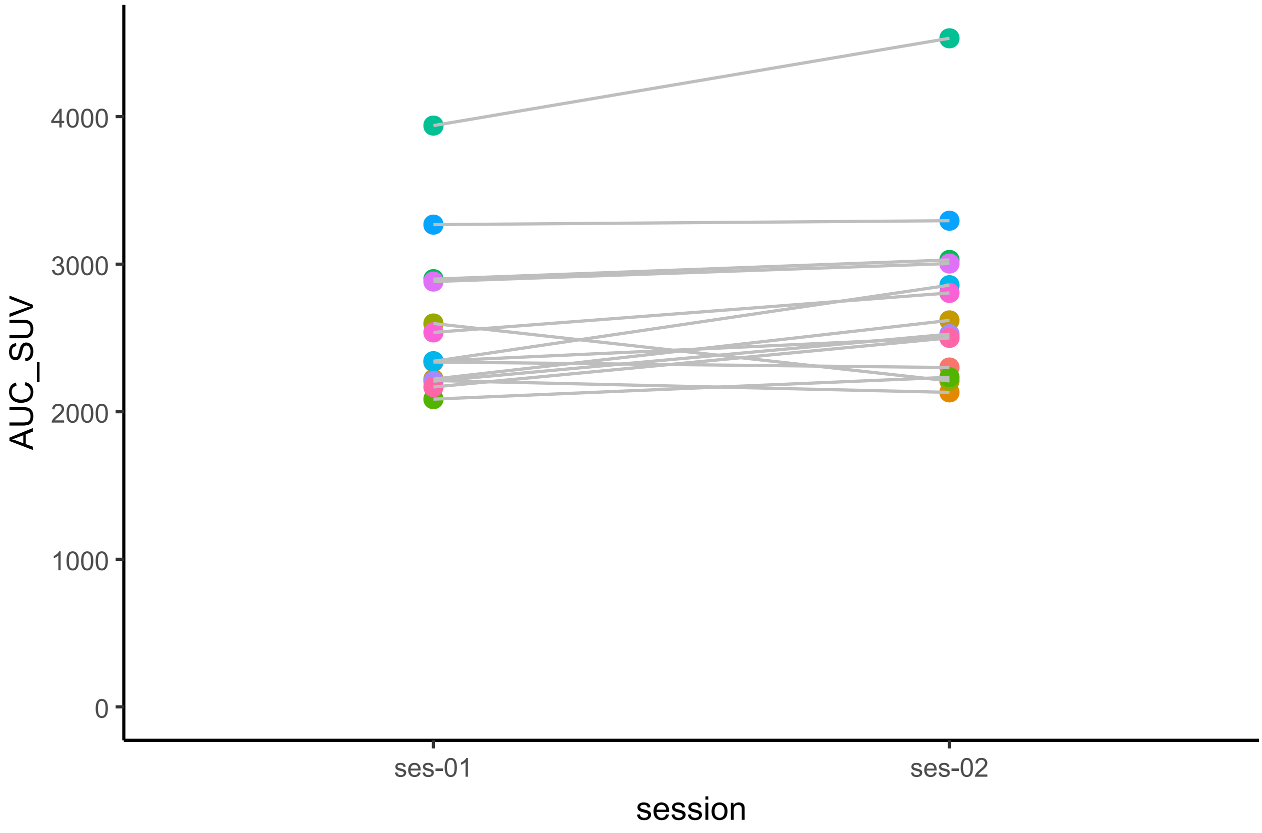


**Figure S11.** Area under the curve of the arterial input function from 0 to 75 min. Data points are color-coded by subject.

**Table S8.** Area under the curve of regional TACs per tool standardized by injected dose (injected radioactivity/body weight) across tools and regions. Note that AUC is calculated with activity in kBq/1000 and time as seconds. Mean difference calculated as ses-02 - ses-01. P-values are calculated using a paired two-tailed t-test.

| Region | ses-01 | ses-02 | Mean difference | p-value |
|---|---|---|---|---|
| | | Solena | | |
| Frontal cortex | 39.649 (5.904) | 40.530 (8.381) | 0.882 | 0.427 |
| Parietal cortex | 40.369 (5.662) | 41.572 (8.328) | 1.202 | 0.288 |
| Temporal cortex | 37.037 (4.710) | 37.905 (6.935) | 0.868 | 0.400 |
| Occipital cortex | 38.140 (4.888) | 39.158 (7.036) | 1.018 | 0.302 |
| Striatum | 40.882 (5.933) | 41.385 (8.233) | 0.503 | 0.658 |
| Hippocampus | 27.280 (3.388) | 27.756 (4.653) | 0.476 | 0.519 |
| Thalamus | 31.870 (4.064) | 32.698 (5.855) | 0.828 | 0.361 |
| Cerebellar cortex | 30.481 (3.662) | 30.983 (4.877) | 0.501 | 0.497 |
| Brainstem | 17.867 (2.274) | 18.328 (3.207) | 0.460 | 0.343 |
| | | PETPrep | | |
| Frontal cortex | 38.620 (5.912) | 39.486 (8.231) | 0.866 | 0.419 |
| Parietal cortex | 39.127 (5.752) | 40.328 (8.222) | 1.200 | 0.263 |
| Temporal cortex | 36.335 (4.793) | 37.137 (6.797) | 0.802 | 0.424 |
| Occipital cortex | 37.908 (5.197) | 38.903 (7.018) | 0.995 | 0.306 |
| Striatum | 39.233 (5.756) | 39.736 (7.930) | 0.504 | 0.642 |
| Hippocampus | 25.398 (3.209) | 25.850 (4.327) | 0.452 | 0.502 |
| Thalamus | 30.037 (3.798) | 30.837 (5.463) | 0.800 | 0.347 |
| Cerebellar cortex | 28.753 (3.518) | 29.230 (4.673) | 0.476 | 0.499 |
| Brainstem | 17.279 (2.072) | 17.688 (2.982) | 0.410 | 0.352 |
| | | PICNIC | | |
| Frontal cortex | 39.934 (5.922) | 40.936 (8.580) | 1.001 | 0.370 |
| Parietal cortex | 40.605 (5.833) | 41.796 (8.456) | 1.191 | 0.292 |
| Temporal cortex | 37.338 (4.735) | 38.225 (7.050) | 0.887 | 0.397 |
| Occipital cortex | 38.565 (5.017) | 39.529 (7.104) | 0.964 | 0.338 |
| Striatum | 41.183 (5.869) | 41.652 (8.179) | 0.469 | 0.680 |
| Hippocampus | 27.358 (3.415) | 27.809 (4.619) | 0.451 | 0.532 |
| Thalamus | 31.882 (4.085) | 32.698 (5.826) | 0.815 | 0.363 |
| Cerebellar cortex | 30.630 (3.596) | 31.107 (4.783) | 0.477 | 0.522 |
| Brainstem | 18.060 (2.238) | 18.522 (3.179) | 0.461 | 0.345 |
